# Optimizing Carbon Sources to Promote Soil Denitrifiers: Lessons for Incubations, Enrichments, and Bioaugmentation

**DOI:** 10.64898/2025.12.18.695110

**Authors:** Louise B. Sennett, Alicia Caro-Pascual, Peter Dörsch, James P. Shapleigh, Åsa Frostegård

## Abstract

Oxygen concentrations fluctuate in soil across time and space. Under anoxic conditions, the three main microbial metabolic pathways - denitrification, fermentation, and DNRA - compete for the same carbon (C) sources. Studies on denitrification in complex soil communities often rely on incubation experiments to determine how various factors affect the regulatory biology of denitrifying organisms and their N_2_O emissions. These experiments typically require an exogenous C source to stimulate measurable activity, and the choice of C source is critical as it should support denitrification while minimizing competition from fermentation and DNRA. This consideration is equally important for the enrichment and isolation of diverse denitrifying organisms and for bioaugmentation-based N_2_O-mitigation strategies. Here, we compared twelve C sources, including glutamic acid, acetate, an artificial root exudate cocktail (eight compounds, individually and in combination), and a clover extract. By combining high-resolution denitrification gas kinetics, metagenomic sequencing, and ^15^N isotope labelling, we aimed to find a C source(s) that (1) supports a diverse soil-derived denitrifying community and (2) limits the competition for C from alternative anaerobic pathways. Among the tested substrates, only the clover extract supported denitrification and maintained a complex denitrifying community. Yet, it also promoted fermentation and DNRA, revealing that a trade-off must exist between fostering denitrifier diversity and limiting growth of organisms using competing anaerobic pathways. More broadly, our results highlight that C source is a methodological fulcrum in controlled soil microbiome studies. It shapes community composition, drives metabolic processes, and ultimately determines the ecological relevance of experimental outcomes and the success of enrichments and soil bioaugmentation approaches.

## Introduction

Unsaturated soils harbour the highest microbial diversity among all environments, driven by the large spatial heterogeneity where cells persist in small, non-competing aggregates [1]. The current knowledge on how various factors influence the composition, activity, and interactions of soil microbiomes is based on studies across a spectrum of complexity, from pure cultures to intact soils. These approaches face an inherent trade-off akin to the Heisenberg uncertainty principle: controlled studies offer better precision but risk introducing artifacts, whereas *in situ* studies better capture natural dynamics but sacrifice experimental precision [2]. The innate heterogeneity of the soil matrix amplifies this trade-off by creating micro-niches with conditions diverging from bulk measurements [3]. To better control the conditions experienced by cells while studying complex communities, a representative fraction of the indigenous population can be extracted from the soil [4] and subjected to ecologically relevant conditions (e.g., fluctuations in O_2_; type and concentration of organic C; NO_3_^-^ concentrations; or pH) in controlled incubations.

Regardless of whether controlled studies on soil microbial processes are performed in microcosms using intact soils or on extracted cells, such investigations generally require an exogenous carbon (C) source to generate measurable microbial activity [5]. The central challenge of these investigations lies in selecting a substrate that sustains the broad microbial diversity inherent to unsaturated soils, while avoiding overgrowth of fast-growing copiotrophs. One ecologically significant soil process driven by organic C is heterotrophic denitrification, as it is governed by a diverse network of facultative anaerobes [6] and is a major biological source of the potent greenhouse gas nitrous oxide (N_2_O) [7]. Dissecting this process under controlled conditions is therefore essential for understanding the regulation of N_2_O in soil communities and guiding mitigation strategies. In this study, we focused specifically on identifying C sources suitable for denitrification research.

During heterotrophic denitrification, microorganisms use organic C as an electron donor to reduce nitrate (NO_3_^-^) to dinitrogen gas (N_2_) via the free intermediates nitrite (NO_2_^-^), nitric oxide (NO), and N_2_O [7]. This process almost always occurs when O_2_ becomes limiting. Exogenous C is typically required in denitrification studies to enable phenotypic measurements or transcription/metatranscriptomic analyses [8–11]. Likewise, it is also needed for the enrichment and/or isolation of diverse denitrifying organisms [12], such as those with a strong N_2_O-reducing capacity [13]. However, the choice of C source poses a major challenge – some may support only narrow subsets of denitrifiers, while others promote competing anaerobic processes, such as fermentation and dissimilatory nitrate reduction to ammonium (DNRA) [14, 15].

Traditionally, soil denitrification studies have oversimplified the choice of C source, with glucose serving as the ‘default’ substrate in a plethora of studies assessing potential denitrification rates or enzyme activities (see long list of references in Morley et al. [16]). This practice persists despite glucose being a poor representation of the low-molecular-weight compounds available to soil microorganisms [16]. Some studies have tested alternative substrates when determining N_2_O reduction rates or N_2_O/N_2_ product ratios [16–19]; however, little effort has been made to identify C sources that preserve community complexity or limit the competition between denitrification, DNRA, and fermentation. The importance of this methodological gap became clear in a recent study: a succinate-based medium, previously recommended for the cultivation of diverse denitrifying organisms from activated sludge [20], yielded only denitrifiers belonging to Pseudomonadales or Rhizobiales when applied to soil-extracted communities [21].

This underscores the need for systematic evaluation of C substrates that support a complex denitrifying community under controlled conditions while limiting competing anaerobic pathways in soil-derived systems to bridge insights on N_2_O regulation from single-organism studies to complex community behavior, to enable the enrichment and isolation of diverse denitrifying organisms with strong N_2_O-reducing capacity, and to understand C competition under hypoxia in soil communities to inform successful bioaugmentation mitigation strategies.

The objective of our study was to: (1) identify C substrates that support a diverse, soil-derived denitrifying bacterial community during anoxic incubation, and (2) assess how these substrates influence the metabolic competition between the main anaerobic processes denitrification, DNRA and fermentation in complex communities, with the goal of stimulating denitrification while minimizing the two other competing pathways. We investigated the effects of 12 C sources on the denitrification phenotypes [11] and composition of a soil-derived bacterial denitrifying community. These C sources included glutamic acid, acetate, an artificial root exudate cocktail containing eight C sources (applied in combination or individually) and a clover leaf extract. The study employed high resolution gas kinetic measurements and metagenomic sequencing to assess microbial changes in community diversity and composition. The results are intended to guide the choice of C-source for various applications in denitrification research, including both intact soils and soil-extracted bacterial communities. Such knowledge is equally important when selecting carrier material for bioaugmentation purposes to ensure desired microbial activity.

## Materials and Methods

### Nycodenz extraction of soil bacteria

To allow precise control over the type of available C substrate, we worked with extracted soil bacterial communities instead of bulk soil. Loamy soil was collected from a grassy ley field in Ås, Norway (59°39’46“N, 10°45’40”E). Field moist soil was sieved (2 mm) and then stored moist (20% gravimetric water content) at 4°C until use. The soil had 3.0% w/w total C content, 0.26% w/w total N content (on a dry weigh [dw] basis), and a pH of 5.2 in a 1:2 0.01M CaCl_2_ slurry.

To raise the bacterial biomass and activity prior to cell extraction, moist soil (25 g fresh weight soil = 20 g dw soil) was mixed with finely ground clover leaves (5 mg dried clover g^-1^ dw soil) and incubated under atmospheric conditions at room temperature for 72 h. Bacterial cells were then extracted from the soil using Nycodenz density gradient centrifugation as described in detail by Bakken and Lindahl [4]. Six parallel Nycodenz extractions were performed from the same soil sample (20 g dw), with three consecutive rounds conducted using the same soil source to obtain sufficient cell numbers. The extracted cells were resuspended and pooled together in a 120 mL sterile vial containing 0.9% NaCl solution and a magnetic stir bar. The vial of pooled cells was incubated and stirred (300 rpm) for ∼24 h in a water bath set at 20°C under atmospheric conditions. Preliminary tests demonstrated that this cell pre-treatment reduced replicate variability and cell aggregation (data not shown). The total number of extracted cells (1.3×10^9^ cells g^-1^ dw soil) was determined by epifluorescence microscopic counts with SYBR green staining [22], representing approximately 20.4% of the total number of cells in the soil.

### Experiment 1

Experiment 1 investigated the effect of different C substrates on denitrification phenotypes and changes in the composition of complex denitrifying communities using amplicon-based and metagenomic sequencing. Four C substrates were tested: acetate, glutamic acid, an artificial root exudate cocktail, and a clover leaf extract. The acetate and glutamic acid treatments were selected as these substrates can be found in soil in the presence of rhizodeposition and cellulose decomposition [18] and play a central role in a wide range of metabolic processes in bacteria [23]. The artificial root exudate cocktail was selected to mimic biologically relevant complex C sources found within the soil rhizosphere. A slightly modified version of the artificial root exudate cocktail described in Paterson et al. [24] was used containing four amino acids (glycine, valine, serine, and alanine) and four organic acids (malic, citric, malonic, and fumaric acid). Sugars were not included to avoid stimulating fermentation. The clover extract was selected to represent a complex C source that is commonly present in agricultural production systems [10, 25]. The preparation and composition of the clover extract is described in detail in the Supplementary Material.

Mineral media were prepared as described by Liu et al. [26] and detailed in the Supplementary Material. The media were prepared using a concentration of 0.3 g C L^-1^ for the sodium acetate, L-glutamic acid, and artificial root exudate treatments. For the artificial root exudate treatments, the concentrations of all compounds were normalized on a C basis, such that 0.0375 g C L^-1^ of each compound was added. Mineral medium without a C source was also prepared for the clover extract treatment. The media were then buffered to pH 6.5 using K_2_HPO_4_ and KH_2_PO_4_ (50 mM), which was selected to mimic the slightly acidic *in situ* conditions of soil while avoiding impaired synthesis of functional N_2_O reductase (NosZ) [10, 26]. The media were dispensed into 120 mL serum vials containing Teflon-coated magnetic stir bars and were then autoclaved. After autoclaving, sterilized (0.2 microns) KNO_3_ was added to each vial, resulting in an initial concentration of 2 mM NO_3_^-^ (100 µmol NO_3_ ^-^ vial^-1^). To the vials containing mineral medium without a C source, 2.35 mL of the sterile filtered clover extract was added. The extract volume was determined through a preliminary experiment to determine the amount of available clover extract needed to provide an amount of available C similar to incubations with 0.3 g C L^-1^ of glutamic acid. This was evaluated by measuring the cumulated production of CO_2_ during an anoxic incubation with 2 mM of NO_3_^-^(data not shown), ensuring sufficient electron donors for complete NO_3_^-^ reduction to N_2_. All vials were then crimp-sealed with butyl septa and made anoxic ([O_2_] ≤10 µM) using an automated manifold by six cycles of gas evacuation and helium (He) filling [27, 28].

The pooled inoculum (0.5 mL) was injected into each crimped-sealed He-washed vial, resulting in a final medium volume of 50 mL and the same initial cell concentration of 1.33×10^9^ cells vial^-1^ (2.66×10^7^ cells mL^-1^) as determined by spectrophotometry (detailed below). Samples from the original inoculum were pelleted and stored at -20°C for DNA extraction and sequencing.

The vials were then placed in a robotic incubation system set at 20°C with a stirring of 800 rpm (detailed below). In total, there were 48 vials divided into four parallel sets to measure all desired parameters. Three sets were used for the continuous headspace measurements of denitrification gas kinetics, NO_3_ ^-^ and nitrite (NO_2_ ^-^) concentrations, and cell numbers throughout the incubation, while a fourth set was used for DNA extraction and sequencing of the bacterial community at the end of the incubation. The clover extract treatment had two additional sets of vials: one designated to continuous measurements of NH_4_^+^ and a second to the measurement of ^15^N-NH_4_^+^ at the end of the incubation (detailed below). All investigated parameters were measured in three biological replicates (n = 3), except for the ^15^N-NH_4_^+^ analysis for which five biological replicates were used (n = 5).

### Experiment 2

Experiment 2 investigated whether all C sources in the artificial root exudate cocktail were utilized as an electron donor by the denitrifying community. Using the same mineral medium described in Experiment 1, the artificial root exudate cocktail was ‘split’ into individual C sources to serve as the sole electron donor per vial (0.3 g C L^-1^). The vials were then adjusted to pH 6.5, autoclaved, amended with NO_3_^-^ (100 µmol NO_3_^-^ vial^-1^), and made anoxic ([O_2_] ≤10 µM) by He-washing, as described in Experiment 1. The soil bacteria were extracted and pooled from the same soil using the same method described in Experiment 1, and 0.5 mL of the pooled inoculum was injected into each vial, resulting in an initial cell concentration of 7.8 x10^7^ cells vial^-1^ (1.56 x10^6^ cells mL^-1^) and a final medium volume of 50 mL. Robot incubation conditions were the same as those described in Experiment 1, in which the headspace of three biological replicates (n=3) per artificial root exudate treatment (24 vials in total) was continuously measured for CO_2,_ N_2_, and N_2_O production to indicate microbial anaerobic respiration on each individual root exudate.

### Experiment 3

Experiment 3 investigated how varying clover extract amendment volumes affected the denitrification phenotypes of complex denitrifying communities. The mineral medium described in Experiment 1 was used, with the concentrations (1x, 1.12x, and 1.22x) adjusted to compensate for the addition of three different clover extract volumes. The pH was adjusted to 6.5, and the vials were autoclaved, amended with NO_3_^-^ (100 µmol NO_3_^-^ vial^-1^) and 2.35 mL, 5.0 mL, or 10 mL vial^-1^ of clover extract, and made anoxic ([O_2_] ≤10 µM) by He-washing as described in Experiment 1. The soil bacteria were extracted and pooled from the same soil using the same method described in Experiment 1, in which 0.5 mL of the pooled inoculum was injected into each vial resulting in 2.09 x10^8^ cells vial^-1^ (4.18 x10^6^ cells mL^-1^) and a final medium volume of 50 mL. Robot incubation conditions were the same as those described in Experiment 1, in which the headspace of three biological replicates (n=3) per clover extract treatment (18 vials in total) was measured frequently for high resolution denitrification gas kinetics.

### Gas kinetics and nitrate/nitrite concentrations

Denitrification kinetics were monitored with frequent headspace measurements of NO, N_2_O, and N_2_ using a robotized incubation system and OpenLAB CDS 2.3 software for gas chromatograph (GC) data acquisition (Agilent), as described by Sennett et al. [11] and detailed further in the Supplementary Material. Samples for NO_3_^-^ and NO_2_^-^ analyses were collected from the liquid (0.1 mL) through the butyl septum of the vials using sterile syringes and analyzed using a chemiluminescence detector system as described by Lim et al. [29] and detailed further in the Supplementary Material.

### Determination of bacterial cell numbers

Cell numbers were determined based on optical density measurements (OD_600_) using the conversion factor 1 OD_600_= 2.0 x10^9^ cells mL^-1^ (determined from microscopic counts with SYBR Green staining) [22]. Cell-specific CO_2_ production rates were calculated using cubic spline interpolation to estimate cell concentration (cells mL^-1^) at the time(s) of CO_2_ rate measurements (mol h^-1^) and expressed as fmol CO_2_ cell^-1^ h^-1^.

### ^15^N enrichment of NH_4_ ^+^

In a preliminary experiment, N-mass balance calculations suggested significant NO_3_ ^-^ reduction via DNRA in the clover extract treatment. To confirm this, a subset of vials in Experiment 1 received a mixed solution of NH_4_NO_3_-^15^N (609455-5G Sigma Aldrich) and KNO_3_, resulting in 2 mM 6.3% ^15^N-NO_3_^-^ per vial. At the end of the incubation, ^15^N-NH_4_^+^ was quantified by chemically converting NH_4_^+^ to N_2_O using the sodium azide (NaN_3_) method, as described in detail by Zhang et al. [30]. Conversion efficiency (98-100%) was confirmed by GC and total NH_4_^+^ pool was measured using a Hach Lange analyzer (DR3900; Ammonium LCK303).

### ^15^N analysis and isotope composition

^15^N abundance in NH_4_^+^ was analyzed by a Finnigan Delta Plus XP isotope ratio mass spectrometer (IRMS) after cryo-focusing and preconcentrating N_2_O in a PreCon unit. Atom percent (At%) of ^15^N was corrected for scale and drift using in-house standards which had been calibrated to the international standards USGS25 and USGS26. To calculate excess ^15^N At% above natural abundance, the At% of unlabeled negative control samples was subtracted from the At% of the labeled samples.

### DNA extraction

After the completion of denitrification (where N_2_ production reached a plateau and N_2_O was completely depleted), untouched vials designated for DNA extraction were spun down and the supernatant was separated from the resulting bacterial pellet. DNA was extracted from the bacterial pellets of the original inoculum and C source treatments using the DNeasy® PowerSoil® Pro kit according to manufacturer protocol (Qiagen, catalog no. 47014). The extracted DNA underwent an additional purification step using the Genomic DNA Clean & Concentrator kit according to the manufacturer protocol (ZymoResearch, catalog no. D4011). The purified DNA was then split into samples for 16S rRNA amplicon-based and metagenomic sequencing (original inoculum and clover extract treatment only) and stored at - 20°C until sequencing.

### ^16^S rRNA amplicon-based and metagenomic sequencing

DNA samples were sent for 16S rRNA gene amplicon and metagenomic sequencing at Novogene (UK) Company Limited in Cambridge, UK. The hypervariable V4 region of the 16S rRNA gene was targeted using the 515F and 806R primers [31] and sequenced using the Illumina NovaSeq 6000 platform (2 × 250 bp paired-end sequencing). The metagenomic sequencing was performed using the Illumina NovaSeq X Plus platform (2 × 150 bp paired-end sequencing). Initial quality control (QC) and library preparation were carried out by the sequencing center, including trimming of sequencing adaptors and barcodes from sequence reads. Detailed bioinformatic workflows and parameters used for the analysis of the 16S rRNA amplicon-based and metagenomic sequence data are provided in the Supplementary Material.

## Results

### Experiment 1

#### Denitrification phenotypes under different C substrates

The glutamic acid treatment strongly supported NO_3_ ^-^ reduction via denitrification **(Fig. 1A)**, depleting NO_3_^-^ by 106 h of incubation, at which N mass balance calculations indicated that 96% of the added NO_3_^-^ was converted to N_2_-N. A transient accumulation of intermediates occurred between 40 and 106 h (max 55.5 µmol NO_2_ ^-^-N, 31.5 nmol NO-N, and 11.8 µmol N_2_O-N per vial) with initial *V*_NAR_, *V*_NIR_, and *V*_NOS_ values of 0.40, 0.17, and 0.12 µmol N vial^-1^ h^-1^, respectively.

**Figure 1.**
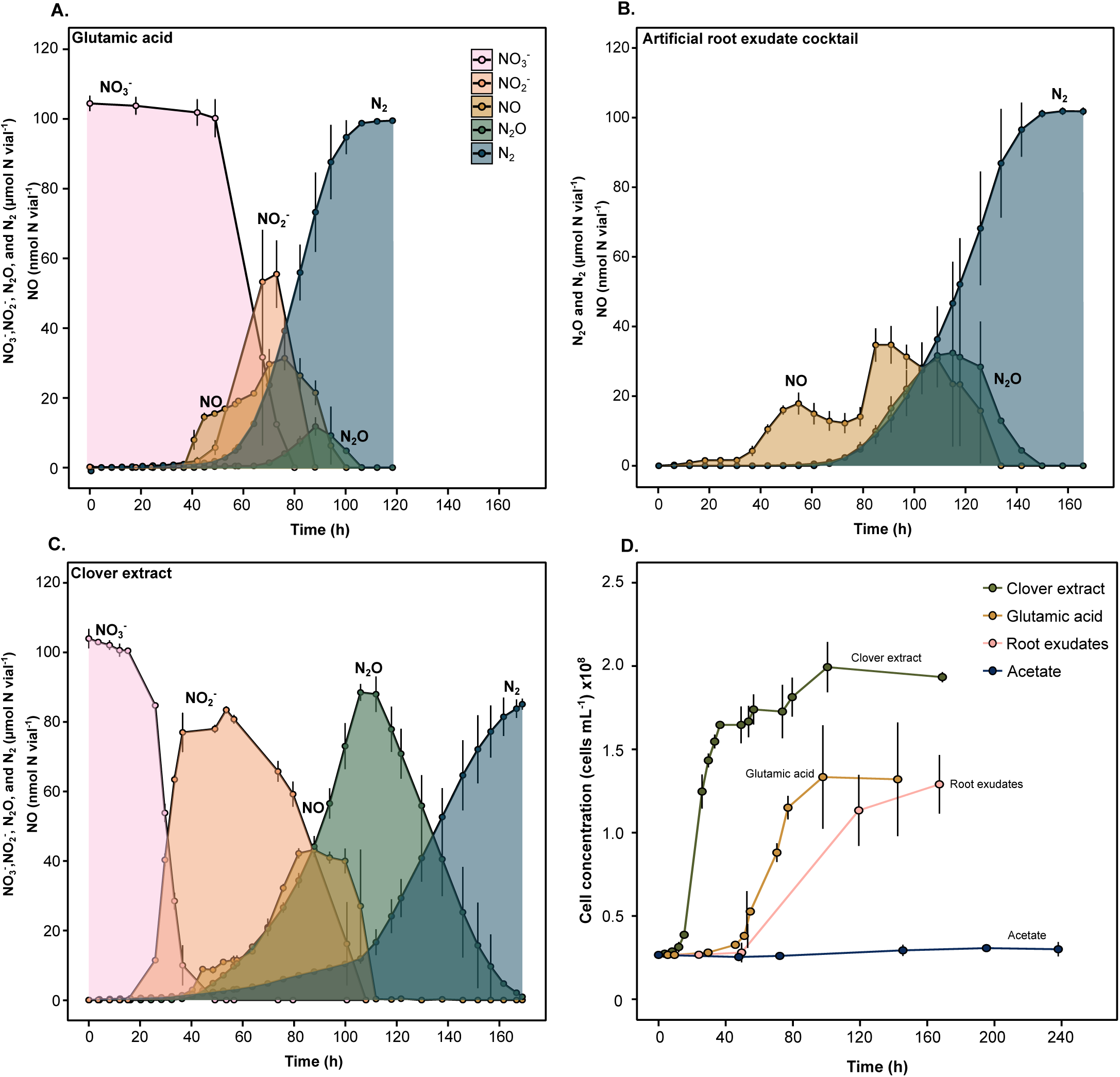
Denitrification phenotypes and cell concentrations of a soil-extracted bacterial community incubated with different carbon substrates. Carbon substrates include (A) glutamic acid, (B) an artificial root exudate cocktail, and (C) clover leaf extract. The denitrification phenotype of the acetate treatment is presented in Supplementary Figure S1. Completion of denitrification was indicated by a plateau in N_2_ in all treatments except the clover extract. In that case, denitrification was considered complete based on the full depletion of NO_3_^-^, NO_2_ ^-^, NO, and N_2_O, despite the absence of a clear N_2_ plateau. (D) Cell concentrations of the bacterial community using glutamic acid, artificial root exudate cocktail, clover extract, or acetate as a carbon substrate. The glutamic acid, root exudate cocktail, and sodium acetate were added at a rate of 0.015 g C vial^-1^ (0.3 g C L^-1^), whereas the clover extract was added at a rate of 2.5 mL vial^-1^ (detailed in Methods). All treatments had an initial cell concentration of 1.33 x10^9^ cells vial^-1^ and contained an initial concentration of 2 mM KNO_3_ (NO_3_^-^ and NO_2_^-^ data are unavailable for the root exudate treatment). The vials were made anoxic prior to the incubation. Data are presented as mean values ± the standard deviation (*n* = 3 biological replicates).

The artificial root exudate cocktail also supported denitrification and did not promote NO_3_^-^ reduction via the DNRA pathway **(Fig. 1B)**, as indicated by the depletion of NO_3_^-^ after 104 h at which N mass balance calculations demonstrated that 100% of the added NO_3_^-^ was converted to N_2_-N. NO production started at 43 h, followed by the simultaneous production of N_2_O and N_2_ after 73 h (NO_3_ ^-^ and NO_2_^-^ data not available). Maximum NO production (34.7 nmol N vial^-1^) was similar to the glutamic acid treatment, while N_2_O production was greater (32.5 µmol N vial^-1^). The artificial root exudate cocktail had an initial *V*_NOS_ of 0.13 µmol N vial^-1^ h^-1^, similar to the glutamic acid treatment (*V*_NAR_ and *V*_NIR_ not available).

The clover extract yielded a distinct denitrification phenotype **(Fig. 1C)**. Nitrate was depleted after 167 h, at which point ^15^N-NH_4_^+^ analysis (0.37 At% excess) indicated that 35% of added ^15^N-NO_3_^-^ was converted to ^15^N-NH_4_^+^ **(Supplementary Table S1).** This suggests that the clover extract treatment significantly promoted NO_3_^-^ reduction via the DNRA pathway. Nitrate reduction started after 15 h, followed by a distinct stepwise reduction of NO_3_^-^ via NO_2_^-^ and N_2_O, during which the peak concentrations of the two intermediates were practically equal to the amount of N finally accumulating as N_2_. The clover extract had a similar initial *V*_NIR_ (0.17 µmol N vial^-1^ h^-1^) compared to the glutamic acid treatment but yielded the greatest initial *V*_NAR_ (3.25 µmol N vial^-1^ h^-1^) and lowest initial *V*_NOS_ (0.06 µmol N vial^-1^ h^-1^) among all treatments that supported denitrification.

In contrast, acetate failed to induce denitrification, as indicated by the absence of NODD reduction during the first 195 h of incubation **(Supplementary Fig. S2)**. After 195 h, there was a slight reduction in NODD, accompanied by the complete depletion of the accumulated NO. This coincided with minor denitrification activity, as evidenced by the production of N_2_O (0.57 µmol N vial^-1^) and N_2_ (2.97 µmol N vial^-1^) after 240 h of incubation.

#### CO_2_ production rates and cell concentrations under different C substrates

The clover extract triggered a burst of CO_2_ production at 20 h, corresponding to a cumulated value of 87 µmol CO_2_ vial^-1^ **(Fig. 2A)** and maximum cell-specific CO_2_ production rate of 0.81 fmol CO_2_ cell^-1^ h^-1^ **(Fig. 2B)**. This was accompanied by an increase in cell concentration from 1.25 x10^8^ to 1.65 x10^8^ cells mL^-1^ **(Fig. 1D)**. At the completion of denitrification, the clover extracted cumulated 143 µmol CO_2_ vial^-1^ and yielded a cell concentration of 1.93 x10^8^ cells mL^-1^.

**Figure 2.**
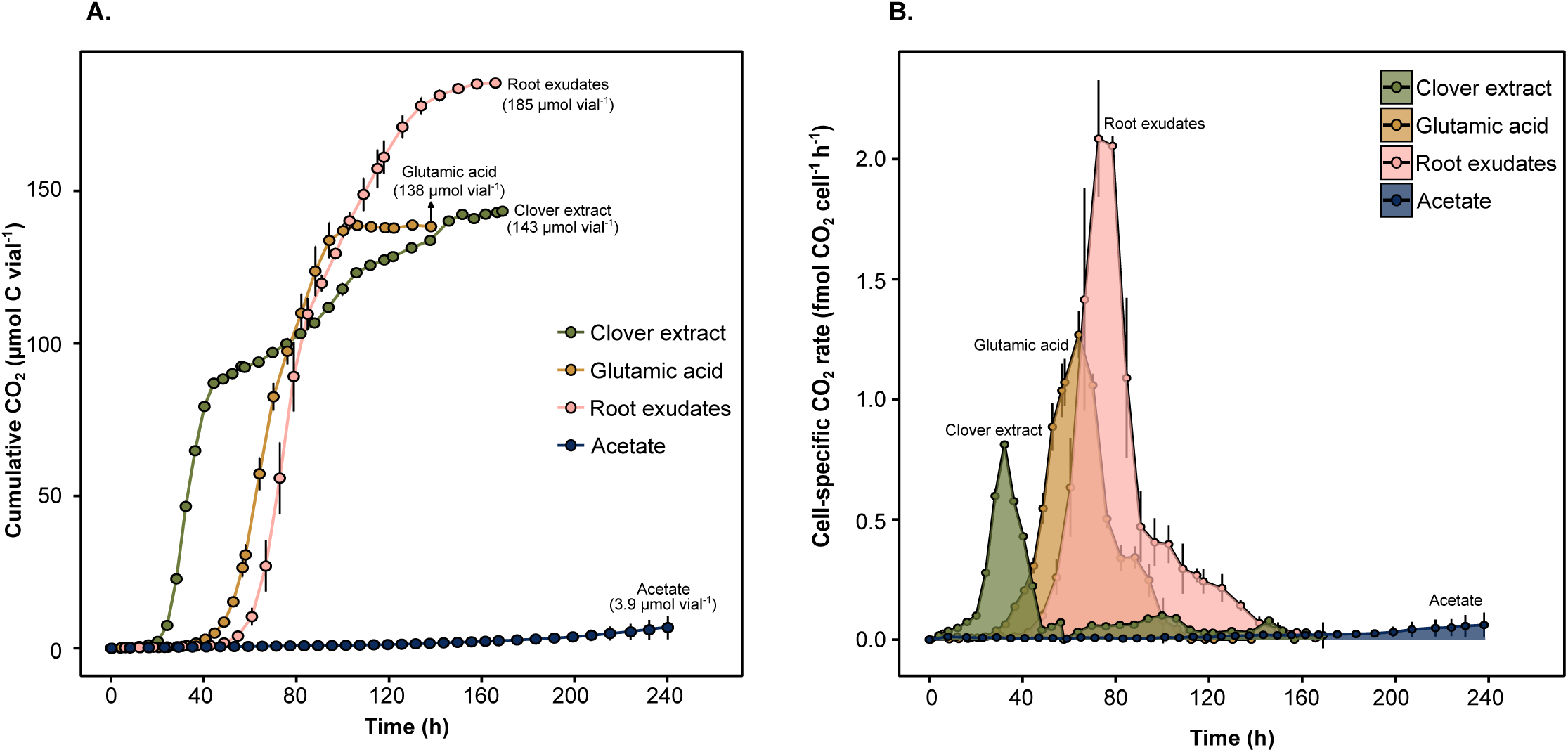
Cumulated CO_2_ and cell-specific CO_2_ production rates of a soil-extracted bacterial community incubated with different carbon substrates under denitrifying conditions. **(A)** cumulated CO_2_ (µmol C vial^-1^) and **(B)** cell-specific CO_2_ production rate (fmol CO_2_ cell^-1^ h^-1^). Carbon substrates included glutamic acid, an artificial root exudate cocktail, clover leaf extract, and sodium acetate. The glutamic acid, root exudate cocktail, and sodium acetate were added at a rate of 0.015 g C vial^-1^ (0.3 g C L^-1^), whereas the clover extract was added at a rate of 2.5 mL vial^-1^ (detailed in Methods). All treatments had an initial cell concentration of 1.33 x10^9^ cells vial^-1^ and contained an initial concentration of 2 mM KNO_3_. The vials were made anoxic prior to the incubation. Data are presented as mean values ± the standard deviation (*n* = 3 biological replicates).

The glutamic acid and artificial root exudate cocktail exhibited similar CO_2_ accumulation patterns, both showing a later burst of CO_2_ production at 40 and 60 h, respectively, with maximum cell-specific CO_2_ production rates of 1.27 and 2.08 fmol CO_2_ cell^-1^ h^-1^, respectively. Final cell concentrations were similar between the treatments (1.32 and 1.29 cells x10^8^ mL^-1^ for glutamic acid and artificial root exudates, respectively), while the artificial root exudate treatment cumulated more CO_2_ than the glutamic acid treatment (185 vs. 138 µmol CO_2_ vial^-1^), likely reflecting alternative anaerobic metabolisms such as fumarate respiration [32].

The acetate treatment exhibited very little CO_2_ production, reaching a maximum cumulated value of 6.8 µmol CO_2_ vial^-1^ after 240 h of incubation **(Fig. 2A)**. This corresponded to very low cell-specific CO_2_ production rates (max 0.06 fmol CO_2_ cell^-1^ h^-1^) and cell growth. Based on these findings, the acetate treatment was omitted from further downstream testing.

#### Bacterial alpha- and beta-diversity under different C substrates

The original inoculum had significantly greater species evenness (0.69 ± 0.01) and richness (884 ± 19.9) compared to all C-substrate treatments (*P* < 0.001), indicating a consistent loss of within-sample diversity regardless of exogenous C substrate **(Fig. 3A).** Among the substrates, the clover extract yielded the greatest Shannon index (1.64 ± 0.03) and species evenness (0.43 ± 0.01), whereas glutamic acid had the greatest richness (64.4 ± 6.72) but the lowest evenness (0.08 ± 0.03). The artificial root exudate treatment had the lowest richness (21.3 ± 2.35).

**Figure 3.**
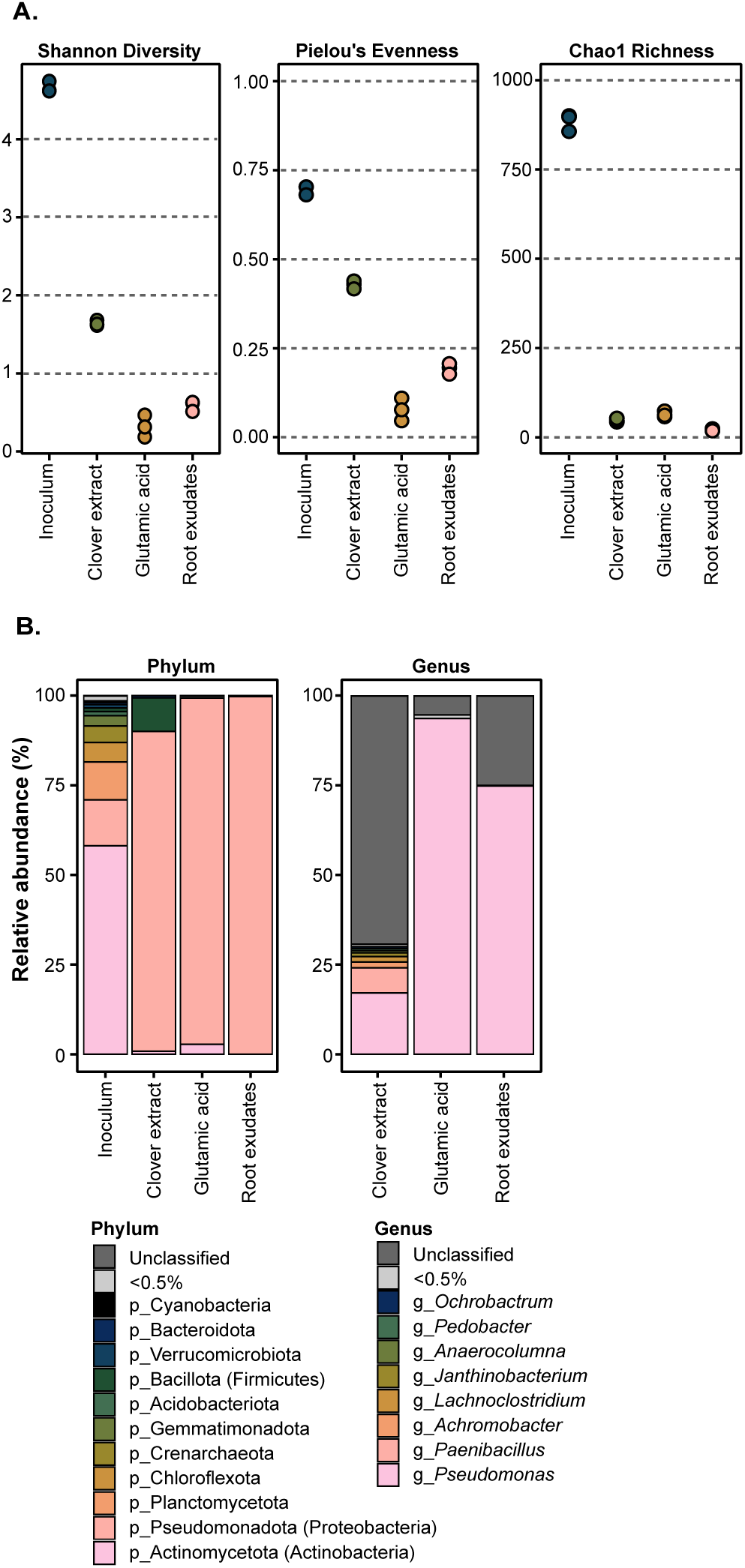
16S rRNA gene-based diversity and taxonomy of a soil-extracted bacterial community incubated with different carbon substrates under denitrifying conditions. **(A)** Alpha-diversity indices (Shannon, Pielou’s evenness, and Chao1 richness) and **(B)** taxonomic assignment at the phylum and genus level. Operational taxonomic units (OTUs) were clustered based on a 97% similarity threshold. For the alpha-diversity indices, samples were rarefied to the minimum sample depth prior to analysis. Treatments include the original inoculum and the bacterial community at the completion of denitrification using either a clover extract, glutamic acid, or an artificial root exudate cocktail as the carbon substrate. Data is presented either as individual biological replicates (diversity indices; n = 3) or as mean values (taxonomy; n = 3).

The ordination of beta-diversity showed a clear separation between the original inoculum and all C substrates along PCoA axis 1 (62.9%) **(Supplementary Fig. S3)**. Axis 2 (27.0%) separated the clover extract from the artificial root exudate and glutamic acid treatments, indicating that the artificial root exudates and glutamic acid treatments had bacterial communities that were compositionally similar to each other but differed from the clover extract treatment.

### Taxonomy of the bacterial community

The original inoculum was mainly populated by Actinomycetota/Actinobacteria (58.1%), followed by Pseudomonadota/Proteobacteria (12.8%), and Planctomycetota (10.6%) **(Fig. 3B)**. Less abundant phyla (< 6%) included Chloroflexota, Crenarchaeota, Gemmatimonadota, Bacillota/Firmicutes, Verrucomicrobiota, Bacteroidota, and Cyanobacteria.

Pseudomonadota/Proteobacteria dominated all C-substrate treatments (89-99.7%). Reads belonging Actinomycetota/Actinobacteria were also present in the glutamic acid (2.81%) and clover extract (0.86%) treatments. Only the clover extract supported additional phyla, including Bacillota/Firmicutes (9.36%) and Bacteroidota (0.59%). At the genus-level, the majority of reads in the clover extract were unassigned (69.3%); among assigned reads, *Pseudomonas* was the most abundant (17.1%), followed by *Paenibacillus* (7.02%) and several low-abundance genera. In contrast, the glutamic acid and artificial root exudate treatments were heavily dominated by *Pseudomonas* (93.7% and 74.9%, respectively).

Overall, the clover extract was the only substrate that supported a relatively complex community, thus downstream analyses using metagenomics focused on its ability to support complexity within the denitrifying fraction of the community.

### Denitrification, DNRA, and fermentation gene abundance and taxonomy

In the clover extract treatment, most denitrification genes (*napA*, *narG*, *nirS*, *cnor*, *nosZ* I) increased in abundance relative to the original inoculum, while *nirK*, *qnor*, and *nosZ* II decreased **(Fig. 4A)**. This suggests that the denitrifying community consisted mainly of canonical denitrifiers [11]. DNRA genes *nirB* and *nirD* also increased, with *nirB* reaching the highest abundance (1110 RPM) **(Fig. 4B)**. The octaheme NO_2_^-^ reductase gene *onr* also increased but remained low in absolute abundance **(inset Fig. 4B).** In contrast, *nrfA* was the only DNRA gene that decreased in abundance **(inset Fig. 4B)**.

**Figure 4.**
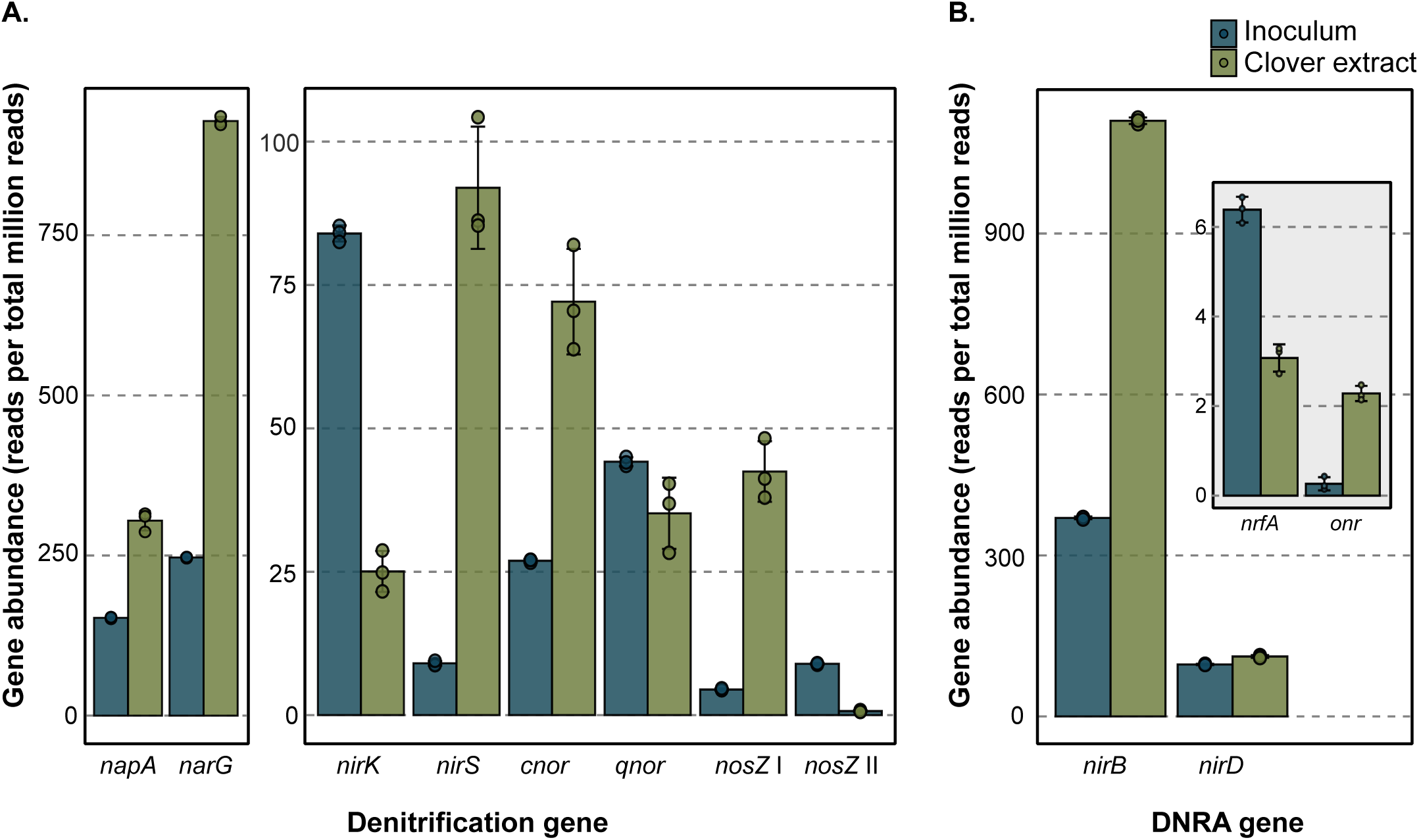
Read abundances of denitrification and DNRA genes from the metagenomes of a soil-extracted bacterial community incubated with clover extract as a carbon source under denitrifying conditions. Abundance of **(A)** denitrification and **(B)** DNRA gene reads. Reads encode NO_3_ ^-^ reductases (*napA* and *narG*), NO_2_ ^-^ reductases (*nirK*, *nirS*, *nrfA*, *nirB*, *nirD*, and *onr*), NO reductases (*cnor* and *qnor*), and N_2_O reductases (*nosZ* clade I and II). Treatments include the original inoculum and the bacterial community at the completion of denitrification using a clover extract as the carbon source. Reads were normalized to reads per total million reads prior to analysis. Data presented as the mean values ± the standard deviation (*n* = 3 biological replicates). The plotted dots represent the individual replicate values.

In the original inoculum, denitrification and DNRA reads were mainly assigned to Actinomycetota/Actinobacteria (*napA*, *narG*, *nirB*, and *nirD*) and Pseudomonadota/Proteobacteria (dominant in *nirK*, *nirS*, *qnor*, *cnor*, *nosZ* I, and *nrfA*) **(Supplementary Fig. S4A)**. *nirK* had notable Thaumarachaeota representation (11%), *nrfA* had notable Verrucomicrobia representation (15%), and *nosZ* II was dominated by Bacteroidota (67.8%).

In the clover extract, 89-99% of denitrification and DNRA reads were assigned to Pseudomonadota/Proteobacteria, except for *nosZ* II, which was dominated by Bacteroidetes (62%), Firmicutes (21%), and Gemmatimonadetes (16%) **(Supplementary Fig. S4B)**. At the genus level, *napA*, *narG*, *nirB*, and *nirD* were populated by *Serratia*, *Pseudomonas*, *Leclercia*, and *Enterobacter*, while *nirK*, *cnor*, *qnor*, and *nosZ* I were populated by *Pseudomonas* and *Achromobacter* **(Fig. 5)**. *nirS* was almost exclusively dominated by *Pseudomonas* (95%). Low-abundance genera (individually < 5%) collectively represented 38 different genera across all denitrification genes **(Supplementary Fig. S5)**, indicating that the clover extract maintained a relatively diverse denitrifying community at the species-level.

**Figure 5.**
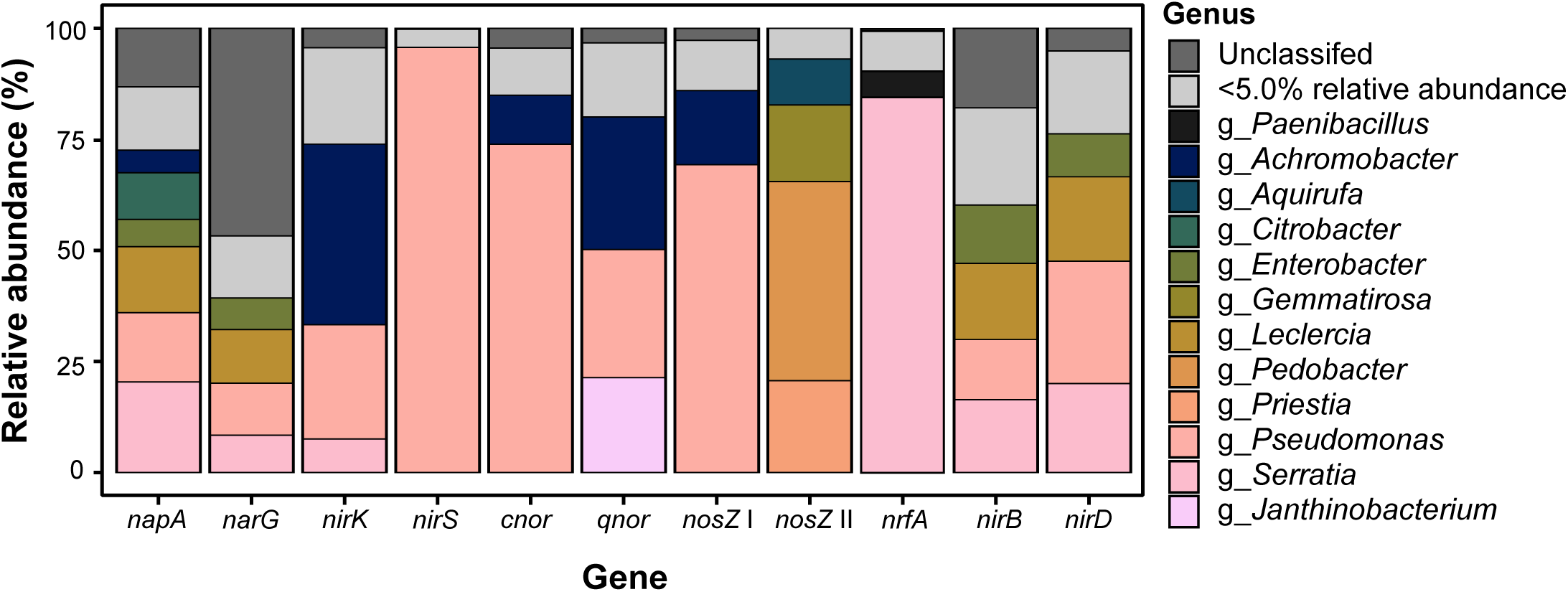
Taxonomic assignment of reads encoding denitrification and DNRA genes from the metagenomes of a soil-extracted bacterial community incubated with clover extract as a carbon substrate under denitrifying conditions. Reads encode NO_3_^-^reductases (*napA* and *narG*), NO_2_^-^ reductases (*nirK*, *nirS*, *nrfA*, *nirB*, and *nirD*), NO reductases (*cnor* and *qnor*), and N_2_O reductases (*nosZ* clade I and II). The data represents gene abundances at the completion of denitrification. Data is presented at the genus level in which genera that individually represented less than 5% of the total relative abundance were grouped together. These are presented individually in Supplementary Figure S4. The data represents the relative abundance presented as the mean values (*n* = 3 biological replicates).

The clover extract increased the abundance of most investigated fermentation-related genes relative to the inoculum, with notable log2 fold-changes in *fumA* (1.47), *mqo* (1.13), *ppc* (1.07), and *pta* (2.16) **(Supplementary Fig. S6)**. These genes were mainly assigned to the genera *Pseudomonas*, *Serratia*, *Leclercia*, and *Enterobacter* **(Supplementary Fig. S7)**.

### Experiment 2

Of the eight C sources in the artificial root exudate cocktail, only fumaric and malic acid were readily utilized as electron donors for denitrification, as shown by the onset of CO_2_ and N_2_ production at ∼60 h of incubation **(Fig. 6A, B)**. Fumaric acid was preferred, as indicated by the earlier completion of denitrification, whereas the malic acid treatment accumulated N_2_O **(Fig. 6C)** and had not reached an N_2_ plateau by the end of the experiment. This suggests differences in the regulatory control of denitrification, depending on C source. The remaining six carbon sources (citric acid, malonic acid, alanine, valine, serine, and glycine) were also used as electron donors for denitrification but at a later stage in the incubation (onset of CO_2_ production at 90-100 h).

**Figure 6.**
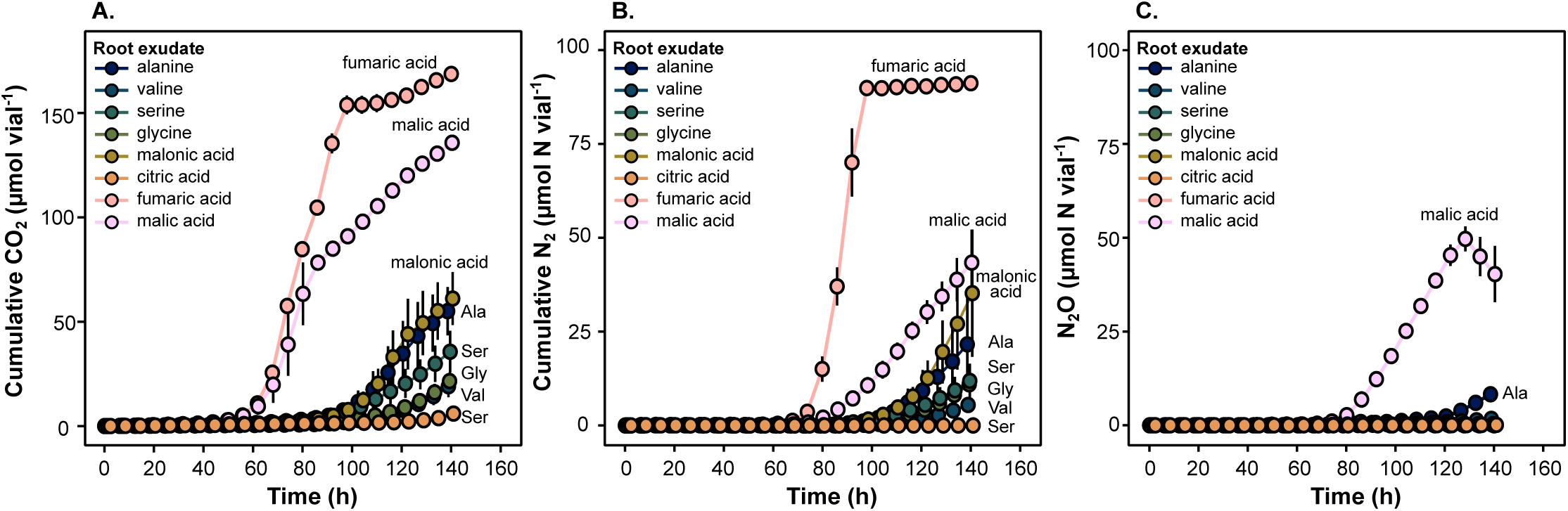
Cumulated CO_2_, cumulated N_2_, and N_2_O production in a soil-extracted bacterial community incubated with different artificial root exudates as a carbon substrate under denitrifying conditions. **(A)** Cumulated CO_2_ (µmol CO_2_ vial^-1^), **(B)** cumulated N_2_ (µmol N_2_-N vial^-1^), and **(C)** N_2_O production (µmol N_2_O-N vial^-1^). The different root exudates included alanine, valine, serine, glycine, malonic acid, citric acid, fumaric acid, and malic acid added at a rate of 0.015 g C vial^-1^ (0.3 g C L^-1^). All treatments had an initial cell concentration of 7.8 x10^7^ cells vial^-1^ and contained an initial concentration of 2 mM KNO_3_ (100 µmol NO ^-^ vial^-1^). The vials were made anoxic prior to the incubation. Data are presented as mean values ± the standard deviation (*n* = 3 biological replicates).

### Experiment 3

Varying volumes of clover extract yielded distinct denitrification phenotypes **(Fig. 7)**. The 5 mL clover treatment produced more CO_2_ (186 µmol vial^-1^) but less N_2_ (65.5 µmol N vial^-1^) and N_2_O (max 57 µmol N vial^-1^) than the 2.35 mL treatment (145 µmol CO_2_, 83 µmol N_2_-N, and 69 µmol N_2_O-N per vial). This pattern intensified at 10 mL, which yielded the most CO_2_ (315 µmol vial^-1^) but the least N_2_ (29 µmol N vial^-1^) and N_2_O (max 19.6 µmol N vial^-1^). Nitrate and nitrite were fully depleted in all treatments.

**Figure 7.**
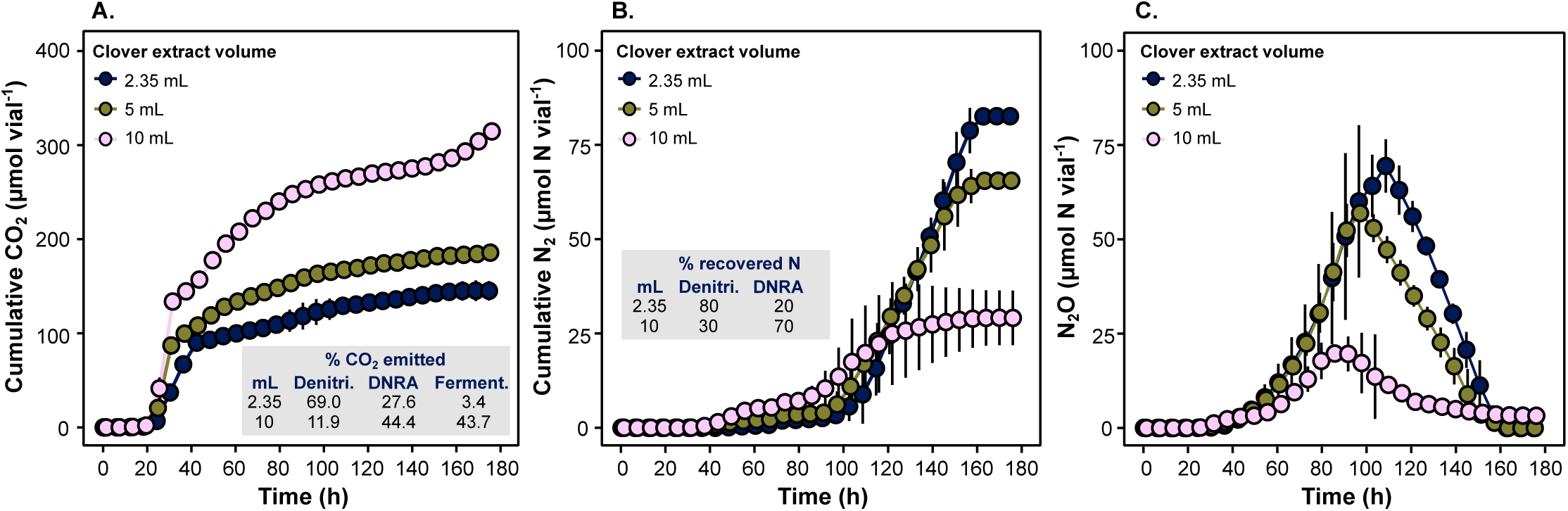
Cumulated CO_2_, cumulated N_2_, and N_2_O production in a soil-extracted bacterial community incubated with different clover extract volumes under denitrifying conditions. **(A)** Cumulated CO_2_ (µmol CO_2_ vial^-1^), **(B)** cumulated N_2_ (µmol N_2_-N vial^-1^), and **(C)** N_2_O production (µmol N_2_O-N vial^-1^). The different clover extract volumes include 2.35 mL, 5 mL, or 10 mL per vial. All treatments had an initial cell concentration of 2.09 x10^8^ cells vial^-1^ and contained an initial concentration of 2 mM KNO_3_ (100 µmol NO ^-^ vial^-1^). The vials were made anoxic prior to the incubation. The percentage of CO_2_ emitted from denitrification (Denitri.), DNRA, and fermentation (Ferment.) are indicated in the inset of panel A, while the percentage of added NO ^-^ recovered N through denitrification and DNRA is indicated in the inset of panel B. Data are presented as mean values ± the standard deviation (*n* = 3 biological replicates). Details for the calculations are provided in the Results.

Electron (*e*-) allocation among denitrification, DNRA, and fermentation was estimated using stoichiometric assumptions: the production of 1 mol CO_2_ from carbohydrates releases 4 mol *e*^-^; the reduction of 1 mol NO_3_^-^ via denitrification (NO_3_^-^ to N_2_) consumes 5 mol *e*^-^; and the reduction of 1 mol NO_3_^-^ via DNRA (NO_3_^-^ to NH_4_^+^) consumes 8 mol *e*^-^ [33]. Using mass balance calculations based on the provided NO_3_ ^-^ and cumulated N_2_-N, 80% of the NO_3_^-^ in the lowest concentration of clover extract was used for denitrification, while 20% was used for DNRA **(inset Fig. 7B)**. These two processes could account for 100 and 40 µmol CO_2_ vial^-1^, respectively, while the remaining 5 µmol CO_2_ (3.4%) was plausibly produced by fermentation **(inset Fig. 7A)**. At the highest extract concentration, 30% of the provided NO_3_ ^-^ was used for denitrification, while 70% was used for DNRA **(inset Fig. 7B)**. These processes could account for 37.5 and 140 µmol of produced CO_2_, respectively. The remaining CO_2_ (44%) was plausibly produced by fermentation **(inset Fig. 7A)**.

## Discussion

Controlled denitrification studies are critical to understanding how N_2_O production is regulated in soil communities and for developing mitigation strategies. Such studies require a C source that supports a diverse denitrifying community, limits competition for C from alternative anaerobic pathways (e.g., DNRA and fermentation), and enables the enrichment/isolation of diverse N_2_O-reducing bacteria [12, 13]. In this study, we evaluated 12 C substrates for their ability to support denitrification in a soil-extracted bacterial community under anoxic incubation. Of these, only one substrate – a clover leaf extract – supported both denitrification and a diverse denitrifying community.

Notably, the clover extract (2.35 mL vial^-1^) was also the only substrate that stimulated DNRA **(Fig. 1C)**, as shown by ^15^N labelling, which revealed ∼35% of the added NO_3_^-^ was converted to NH_4_^+^. This aligns with our previous study, where N-mass balance calculations indicated that 11-23% of the added NO_3_^-^ went to DNRA in soil amended with clover residues [11]. Stoichiometric calculations also demonstrated that ∼3% of the produced CO_2_ in the low concentration clover extract treatment originated from fermentation, which coincided with an observed increase in several fermentation-related genes **(Supplementary Fig. S6)**. The co-occurrence of DNRA and fermentation in this treatment contrasts our original notion of what constitutes an optimal C source for denitrification studies and highlight the core challenge in selecting a substrate that selectively enriches for a community of diverse denitrifying organisms without promoting competing anaerobic pathways.

In slow-growing systems, partial denitrifiers often outnumber complete denitrifiers [12, 34]. Many organisms that carry one denitrification gene are capable of performing DNRA, fermentation, and other metabolic processes involving NO_2_^-^ [35–37]. Therefore, to accurately capture the functional and taxonomic diversity of denitrifiers in controlled incubation systems, while avoiding the overgrowth of ‘weedy organisms’, it may be necessary to tolerate the co-enrichment of organisms performing alternative anaerobic metabolisms. This represents a trade-off between targeted enrichment and microbial diversity in denitrification incubation studies.

Although gene abundance does not directly reflect microbial activity, the marked increase in *nirB* and decrease in *nrfA* **(Fig. 4B)** indicate that DNRA in the clover extract treatment was primarily fermentative rather than respiratory [38, 39]. This is consistent with the enrichment *nirB*-containing organisms belonging to *Enterobacter*, *Serratia*, *Pseudomonas*, and *Leclercia* **(Fig. 5)**, which were also associated with the increase of fermentation-related genes **(Supplementary Fig. S7)**. It is widely accepted that DNRA is favoured over denitrification under NO_3_^-^ limiting conditions with excess electron donor [15, 33, 40, 41]. However, it has also been proposed that highly reduced fermentable C sources may further promote fermentative DNRA, as these organisms can derive energy from both substate-level phosphorylation and electron donor oxidation [42]. This indicates that C source quality – not just the C/NO_3_ ^-^ ratio of available substates – influences the balance of DNRA and denitrification in complex communities.

The NO_3_^-^ and NO_2_ ^-^ kinetics suggest that fermentative DNRA dominated in the clover extract treatment during the first ∼40 h **(Fig. 1C)**, consistent with the elevated CO_2_ production observed at this time **(Fig. 2B)**. DNRA later lost its competitive edge over denitrification, either due to a declining C/NO_3_^-^ ratio of available substrates, shifting electron flow to denitrification [14], or a preferential depletion of specific C sources via fermentation or DNRA. These ideas are both supported by increased clover extract volumes leading to reduced amounts of NO_3_^-^ recovered as N_2_ **(Fig. 7B)** and increased proportions of fermentation-derived CO_2_ **(Fig. 7A)**. This competition for C and NO_3_^-^ may explain why the clover extract uniquely supported a diverse community [43, 44].

Both the glutamic acid and artificial root exudate treatment exhibited a ∼20 h delay in NO_3_^-^ reduction **(Fig. 1A)**, cell growth **(Fig. 1D)**, and CO_2_ production **(Fig. 2)**, likely due to the initially low abundance of denitrifiers able to use these substates. Both treatments were dominated by *Pseudomonas* **(Fig. 3B)**, particularly the glutamic acid treatment (98% relative abundance). In our study, this dominance may represent a ‘hot moment’ in soil [45], where a sudden availability of labile C (depending on its quality) triggers rapid denitrification activity on a localized microscale by organisms that are well-adapted to capitalize on transient pulses of resources. Notably, the glutamic acid treatment also had the greatest denitrification rates, thus appearing as a promising C substrate for community studies; however, this activity originated from a very narrow subset of denitrifying organisms. This finding highlights the importance of pairing community data with detailed phenotyping in order to avoid biased interpretations.

Despite providing a ‘complex’ C substrate, the artificial root exudate cocktail still yielded a community dominated (75%) by *Pseudomonas* **(Fig. 3B)**. This, however, is consistent with diversity being lower in the rhizosphere compared to bulk soil due to the dominance of fast-growing copiotrophs [46]. When divided into individual C substrates (Experiment 2), it was evident that only fumaric and malic acid were readily used for denitrification **(Fig. 6)**, which likely persisted when the exudates were applied in combination (Experiment 1), as the patterns for CO_2_, N_2_O, and N_2_ production closely resembled each other in both experiments **(Figs. 1B and 2A)**. Preference for these substrates may reflect a C catabolite repression mechanism that has been documented in *Pseudomonas spp*. [47–50], enabling selective assimilation of preferred C sources that improves their competitiveness in natural habitats [50].

The acetate treatment was the only C substrate that did not support denitrification or microbial activity in our soil-extracted bacterial community, as judged from the absence of N_2_ production **(Supplementary Fig. S2)**, CO_2_ production **(Fig. 2A)**, or cellular growth **(Fig. 1D)** after 240 h of incubation. This was surprising, as acetate is a commonly used C substrate for denitrification in wastewater treatment plants [51]. The apparent lack of microbial response to acetate in our study suggests that this particular bacterial community was not adapted to using acetate as a C source in its native environment, raising the possibility that assimilation may vary across soil communities. This aligns with a previous study that demonstrated that acetate assimilation was not universal among all denitrifying bacteria in an activated sludge system [52].

Although the clover extract, to some extent, promoted both DNRA and fermentation, it was the only substrate out of 12 that supported a diverse denitrifying community. These findings suggest that a trade-off must be tolerated between fostering denitrifier diversity and limiting growth of organisms using competing anaerobic pathways. More broadly, our results underscore that the choice of C source is a methodological fulcrum in controlled soil microbiome studies, determining which organisms thrive and metabolic processes dominate.

Recognizing this is essential for designing experiments that preserve the ecological complexity of soil microbial communities and generate meaningful results. Finally, lessons learned from approaches presented in the present study will help guide the development of suitable carrier materials for bioaugmentation with N_2_O reducing bacteria [13], supporting the inoculant while avoiding the stimulation of organisms performing other metabolisms and competing for the same C source.

## Author contributions

L.B.S. and Å.F. designed the experiments. L.B.S. and A.C.P. collected the data. L.B.S., A.C.P., P.D., and J.P.S. analyzed the data. All authors contributed to the interpretation of results and writing of the manuscript.

## Acknowledgments

We thank Prof. Lars R. Bakken for his valuable input towards the interpretation of our results and Lars Molstad for his contribution to the concept of Experiment 2.

## Data availability

The 16S rRNA amplicon and metagenomic sequence data generated in this study have been deposited as FASTQ files in the European Nucleotide Archive (ENA) under BioProject PRJEB90840.

## References

1. Zhou J, Xia B, Treves DS et al. Spatial and resource factors influencing high microbial diversity in soil. Appl Environ Micobiol 2002;68:326–34. 10.1128/AEM.68.1.326–334.2002

2. Madsen E. Identifying microorganisms responsible for ecologically significant biogeochemical processes. Nat Rev Microbiol 2005;3:439–46. 10.1038/nrmicro1151

3. Wilpiszeski RL, Aufrecht JA, Retterer ST et al. Soil aggregate microbial communities: Towards understanding microbiome interactions at biologically relevant scales. Appl Environ Micobiol 2019;85:e00324–19. 10.1128/AEM.00324-19.

4. Bakken L, Lindahl V. Recovery of bacterial cells from soil. In: Elsas Jv, Trevors J (eds.), Nucelic acids in the environment: Methods and applications, Springer. 1995, 9–27.

5. Anderson JPE, Domsch KH. A physiological method for the quantitative measurement of microbial biomass in soils. Soil Biol Biochem 1978;10:215–21.

6. De Graf RH, Jones CM, Hallin S. Intergenomic comparisons highlight modularity of the denitrification pathway and underpin the importance of community structure for N_2_O emissions. PLoS ONE 2014;9:e114118. 10.1371/journal.pone.0114118

7. Zumft WG. Cell biology and molecular basis of denitrification. Microbiol Mol Biol Rev 1997;61:533–616.

8. Henderson SL, Dandie CE, Patten CL et al. Changes in denitrifier abundance, denitrification gene mrna levels, nitrous oxide emissions, and denitrification in anoxic soil microcosms amended with glucose and plant residues. Appl Environ Micobiol 2010;76:2155–64. 10.1128/AEM.02993-09

9. Wertz S, Goyer C, Zebarth BJ et al. Effects of temperatures near the freezing point on N_2_O emissions, denitrification and on the abundance and structure of nitrifying and denitrifying soil communities. FEMS Microbiol Ecol 2013;83:242–54. 10.1111/j.1574-6941.2012.01468.x

10. Frostegård Å, Vick SHW, Lim NYN et al. Linking meta-omics to the kinetics of denitrification intermediates reveals ph-dependent causes of N_2_O emissions and nitrite accumulation in soil. ISME J 2022;16:26–37. 10.1038/s41396-021-01045-2

11. Sennett LB, Roco CA, Lim NYN et al. Determining how oxygen legacy affects trajectories of soil denitrifier community dynamics and N_2_O emissions. Nat Commun 2024;15 10.1038/s41467-024-51688-w

12. Lycus P, Bøthun KL, Bergaust L et al. Phenotypic and genotypic richness of denitrifiers revealed by a novel isolation strategy. ISME J 2017;11:2219–32. 10.1038/ismej.2017.82

13. Hiis EG, Vick SHW, Molstad L et al. Unlocking bacterial potential to reduce farmland N_2_O emissions. Nature 2024;630:421–28.

14. van der Berg E, Elisário MP, Kuenen JG et al. Fermentative bacteria influence the competition between denitrifiers and DNRA bacteria. Front Microbiol 2017;8:1684. 10.3389/fmicb.2017.01684

15. Kraft B, Tegetmeyer H, Sharma R et al. The environmental controls that govern the end product of bacterial nitrate respiration. Science 2014;345:676–79. 10.1126/science.1254070

16. Morley NJ, Richardson DJ, Baggs EM. Substrate induced denitrification over or under estimates shifts in soil N_2_/N_2_O ratios. PLoS ONE 2014;9:e108144. 10.1371/journal.pone.0108144

17. Murray PJ, Hatch DJ, Dixon ER et al. Denitrification potential in a grassland subsoil: Effect of carbon substrates. Soil Biol Biochem 2004;36:545–47. 10.1016/j.soilbio.2003.10.020

18. Morley N, Baggs EM. Carbon and oxygen controls on N_2_O and N_2_ production during nitrate reduction. Soil Biol Biochem 2010;42:1864–71. 10.1016/j.soilbio.2010.07.008

19. Maheshwari A, Jones CM, Tiemann M et al. Carbon substrate selects for different lineages of N_2_O reducing communities in soils under anoxic conditions. Soil Biol Biochem 2023;177:108909. 10.1016/j.soilbio.2022.108909

20. Heylen K, Vanparys B, Wittebolle L, et al. Cultivation of denitrifying bacteria: Optimization of isolation conditions and diversity study. Appl Environ Micobiol 2006;72:2637–43. 10.1128/AEM.72.4.2637-2643.2006

21. Crocker K, Lee KK, Chakraverti-Wuerthwein M et al. Environmentally dependent interactions shape patterns in gene content across natural microbiomes. Nature Microbiology 2024;9:2022–37. 10.1038/s41564-024-01752-4

22. Noble R, Fuhrman J. Use of sybr green i for rapid epifluorescence counts of marine viruses and bacteria. Aquat Microb Ecol 1998;14:113–18.

23. Berg JM, Tymoczko JL, Stryer L. Biochemistry 6edn. New York, NY: Friedman & Co., 2007.

24. Paterson E, Gebbing T, Abel C et al. Rhizodeposition shapes rhizosphere microbial community structure in organic soil. New Phytologist 2007;173:600–10. 10.1111/j.1469-8137.2006.01931.x

25. Liu B, Mørkved PT, Frostegård Å et al. Denitrification gene pools, transcription and kinetics of NO, N_2_O and N_2_ production as affected by soil pH. FEMS Microbiol Ecol 2010;72:407–17. 10.1111/j.1574-6941.2010.00856.x

26. Liu B, Frostegård Å, Bakken LR. Impaired reduction of N_2_O to N_2_ in acid soils is due to a posttranscriptional interference with the expression of *nosz*. mBio 2014;5:e01383–14. 10.1128/mBio.01383-14.

27. Molstad L, Dörsch P, Bakken LR. Robotized incubation system for monitoring gases (O_2_, NO, N_2_O, N_2_) in denitrifying cultures. J Microbiol Methods 2007;71:202–11. 10.1016/j.mimet.2007.08.011

28. Molstad L, Dörsch P, Bakken L Improved robotized incubation system for gas kinetics in batch cultures. Researchgate. 10.13140/RG.2.2.30688.07680.

29. Lim NYN, Frostegård Å, Bakken LR. Nitrite kinetics during anoxia: The role of abiotic reactions versus microbial reduction. Soil Biol Biochem 2018;119:203–09. 10.1016/j.soilbio.2018.01.006

30. Zhang L, Altabet MA, Wu T et al. Sensitive measurement of NH_4_^+^ ^15^N/^14^n(δ^15^NH_4_^+^) at natural abundance levels in fresh and saltwaters. Anal Chem 2007;79:5297–303. 10.1021/ac070106d

31. Caporaso JG, Lauber CL, Walters WA et al. Global patterns of 16s rRNA diversity at a depth of millions of sequences per sample. Proc Natl Acad Sci USA 2011;108:4516–22. 10.1073/pnas.1000080107

32. Unden G, Bongaerts J. Alternative respiratory pathways of *escherichia coli*: Energetics and transcriptional regulation in response to electron acceptors. Biochimica et Biophysica Acta (BBA) - Bioenergetics 1997;1320:217–34. 10.1016/S0005-2728(97)00034-0

33. Tiedje JM, Sextone AJ, Myrold DD et al. Denitrification: Ecological niches, competition and survival Antonie van leeuwenhoek 1982;48:569-83. 10.1007/BF00399542

34. Roothans N, van Loosdrecht MC, Laureni M. Metabolic labour division trade-offs in denitrifying microbiomes. ISME J 2025;19:wraf020. 10.1093/ismejo/wraf020

35. Sanford RA, Wagner DD, Wu Q et al. Unexpected nondenitrifier nitrous oxide reductase gene diversity and abundance in soils. PNAS 2012;109:19709–14. 10.1073/pnas.1211238109

36. 4Mania D, Heylen K, Spanning RJMv et al. Regulation of nitrogen metabolism in the nitrate ammonifying soil bacterium *Bacillus vireti* and evidence for its ability to grow using N_2_O as electron acceptor. Environ Microbiol 2016;18:2937–50. 10.1111/1462-2920.13124

37. Zhang IH, Sun X, Jayakumar A et al. Partitioning of the denitrification pathway and other nitrite metabolisms within global oxygen deficient zones. ISME Communications 2023;3:76. 10.1038/s43705-023-00284-y

38. Pandey CB, Kumar U, Kaviraj M et al. DNRA: A short-circuit in biological N-cycling to conserve nitrogen in terrestrial ecosystems. Sci Total Environ 2020;788:139710. 10.1016/j.scitotenv.2020.139710

39. Kaviraj M, Kumar U, Snigdha A et al. Nitrate reduction to ammonium: A phylogenetic, physiological, and genetic aspects in prokaryotes and eukaryotes. Archives of Microbiology 2024;206:297. 10.1007/s00203-024-04009-0

40. Rütting T, P Boeckx, Müller C et al. Assessment of the importance of dissimilatory nitrate reduction to ammonium for the terrestrial nitrogen cycle. Biogeosciences 2011;8:1779–91. 10.5194/bg-8-1779-2011

41. Yoon S, Cruz-Garcia C, Sanford R et al. Denitrification versus respiratory ammonification: Environmental controls of two competing dissimilatory NO_3_^-^ /NO_2_^-^ reduction pathways in *Shewanella loihica* strain PV-4. ISME J 2015;9:1093–104. 10.1038/ismej.2014.201

42. Rehr B, Klemme J. Competition for nitrate between dnitrifying *Pseudomonas stutzeri* and nitrate ammonifying enterobacteria. FEMS Micrbiol Ecol 1989;62:51–58.

43. Bello MD, Lee H, Goyal A et al. Resource–diversity relationships in bacterial communities reflect the network structure of microbial metabolism. Nature Ecology & Evolution 2021;5:1424–34. 10.1038/s41559-021-01535-8

44. Lee H, Bloxham B, Gore J. Resource competition can explain simplicity in microbial community assembly. PNAS 2023;120:e2212113120. 10.1073/pnas.2212113120

45. McClain ME, Boyer EW, Dent CL et al. Biogeochemical hot spots and hot moments at the interface of terrestrial and aquatic ecosystems. Ecosystems 2003;6:301–12. 10.1007/s10021-003-0161-9

46. Philippot L, Raaijmakers JM, Lemanceau P et al. Going back to the roots: The microbial ecology of the rhizosphere. Nature Reviews in Microbiology 2013;11:789–99. 10.1038/nrmicro3109

47. Hester K, Madhusudhan K, Sokatch J. Catabolite repression control by *crc* in 2xyt medium is mediated by posttranscriptional regulation of *bkdr* expression in *Pseudomonas putida*. J Bacteriol 2000;182:1150–53. 10.1128/jb.182.4.1150-1153.2000

48. Morales G, Linares J, Beloso A et al. The *Pseudomonas putida* crc global regulator controls the expression of genes from several chromosomal catabolic pathways for aromatic compounds. J Bacteriol 2004;186:1337–44. 10.1128/jb.186.5.1337-1344.2004

49. Moreno R, Martınez-Gomariz M, Yuste L et al. The *Pseudomonas putida* crc global regulator controls the hierarchical assimilation of amino acids in a complete medium: Evidence from proteomic and genomic analyses. Proteomics 2009;9:2910–28. 10.1002/pmic.200800918

50. Rojo F. Carbon catabolite repression in Pseudomonas: Optimizing metabolic versatility and interactionswith the environment. FEMS Microbiol Rev 2010;34:658–84. 10.1111/j.1574-6976.2010.00218.x

51. Henze M, van Loosdrecht MC, Brdjanovic D. Biological wastewater treatment. IWA publishing, 2008. 10.2166/9781780401867

52. Thomsen TR, Kong Y, Nielsen PH. Ecophysiology of abundant denitrifying bacteria in activated sludge. FEMS Micrbiol Ecol 2007;60:370–82. DOI:10.1111/j.1574-6941.2007.00309.x

