## Supplementary Material for "Optimizing Carbon Sources to Promote Soil Denitrifiers: Lessons for Incubations, Enrichments, and Bioaugmentation"

**Content:**

Supplementary Materials and Methods

Supplementary Table S1 and S2

Supplementary Figures S1-7

References

**Supplementary Materials and Methods**

**Preparation of the clover leaf extract**

The clover extract was prepared using a rate of 20 mg dried red clover leaf powder mL^-1^ dH_2_O. The suspension was brought to a boil in a microwave and then placed on a magnetic plate stirring continuously and heated to 50°C for 2 h. The suspension was then vacuum filtered to remove the clover residues and the resulting extract was sterile filtered (0.2 microns) and stored at 4°C until use. The clover extract was analyzed for its polysaccharide composition using ion chromatography, employing both mono- and oligosaccharide detection methods [1]. This screening suggested that the extract contained high concentrations of simple sugars – primarily sucrose, glucose, and fructose – along with lower levels of longer, unidentified oligosaccharides.

**Mineral Medium**

The medium was the same as that previously described by Liu et al. [2] with the exception that 6 mM rather than 3 mM of (NH_4_)_2_SO_4_ was used to prevent NO_3_^-^ assimilation: 6 mM(NH_4_)_2_SO_4_, 1.76 mg L^-1^ EDTA, 10 mg L^-1^ ZnSO4, 5 mg L^-1^ FeSO_4_, 1.5 mg L^-1^ MnSO_4_, 0.4 mg L^-1^ CuSO_4_, 0.25 mg L^-1^ Co(NO_3_)_2_, 0.15 mg L^-1^ H_3_BO_3_, 1 mg L^-1^ nicotinic acid, 0.5 mg L^-1^ thiamine, and 1 mg L^-1^ biotin, buffered to pH 6.5 using 50 mM phosphate buffer (K_2_HPO_4_ and KH_2_PO_4_)

**Gas kinetics and robotic incubation system**

Denitrification kinetics were monitored with frequent headspace measurements of NO, N_2_O, and N_2_ using a robotized incubation system and OpenLAB CDS 2.3 software for gas chromatograph (GC) data acquisition (Agilent), as described in detail by Molstad et al. [3, 4]. Briefly, the vials were maintained in a thermostatic water bath. Gas samples were taken at intervals from the headspace by an autosampler coupled to a peristaltic pump. Gas samples were pumped into a gas chromatograph (Varian; 7890A GC, Agilent) for analysis of CO_2_, O_2_, N_2_O and N_2_, and into a chemiluminescence NOx analyzer (M200E, Teledyne) for analysis of NO. For each sampling, an equal volume of He was pumped back into the vial to maintain ~1 atm pressure. Measurements for N_2_ were corrected for dilution and contamination with atmospheric N that took place when sampling the headspace, as described in Molstad et al. [3]

**Nitrate and Nitrite Concentrations**

Samples for NO_3_^-^ and NO_2_^-^ analysis were collected from the liquid (0.1 mL) through the butyl septum of the vials using sterile syringes and analyzed as described by Lim et al. [5]. Briefly, 10 µL of the sample was immediately injected into a purging device with reducing agents: either acetate+NaI (at room temperature) for converting NO_2_^-^ to NO, or HCl+VCl_3_ (at 95 ^0^C) for converting NO_3_^-^+NO_2_^-^ to NO. The NO was transported by a stream of N_2_ to a chemiluminescence detector system. The integrated NO peaks were used to estimate the concentration of NO_2_^-^ or NO_3_^-^+NO_2_^-^, in which NO_3_^-^ concentrations were determined by subtracting the NO_2_^-^ from the NO_3_^-^+NO_2_^-^ values.

**Analysis of 16S rRNA amplicon-based sequence data**

Amplicon sequence data was processed using the QIIME2 pipeline (v.2021.8) [6]. The sequences were demultiplexed and then denoised and trimmed using the DADA2 plugin [7] where only reads in the length range of 221-223 bp were retained. Trimmed forward and reverse reads were then merged and only successfully merged reads were retained. Clustering of operational taxonomic units (OTUs) was performed at 97% sequence similarity. Representative sequences from each OTU were classified by finding their closest match in a set of reference 16S rRNA gene sequences and using the Naïve Bayes Classifier trained on SILVAs 16S rRNA gene sequences of the 515F/806R region (v.138) [8]. The relative abundance of taxonomic groups found within the bacterial communities was performed using Microbiome Analyst [9, 10] with no additional filters. Bacterial beta-diversity was assessed using the vegan package (v.2.6-6.1) [2] in RStudio (v.4.4.1). Data were square root transformed, and Bray-Curtis dissimilarities were generated. Differences in bacterial community composition among treatments were ordinated using a principal coordinate analysis (PCoA) plot generated with ggplot2 (v.3.5.1) in RStudio. Samples were rarefied to 38438 reads per sample for the purpose of determining bacterial alpha diversity indices (Shannon index, Chao1 species richness, and Pielou’s evenness) using the vegan package in RStudio.

**Analysis of metagenomic sequence data**

Read quality was verified using FastQC (v.0.12.1) in KBase [11]. The paired-end reads were then trimmed using Trimmomatic (v.0.36) [12] in KBase. After trimming, the number of reads (in millions) was between 102.5 and 150.2 for the original inoculum and between 79.7 and 82.1 for the clover extract treatment. For functional annotation, reads were aligned using DIAMOND [13] with an e-value cut-off of 1x10^-3^ and compared against protein reference data sets retrieved from AnnoTree [14] for genes involved in denitrification (*narG*/*napA*, *nirK*/*nirS*, and *cnor*/*qnor*), other NO_3_^-^ consumption pathways (DNRA) (*nrfA*, *nirB*, and *nirD*) [15], and fermentation-related genes based on KEGG annotations (*ackA*, *acs*, *adh*, *E1.1.1.90*, *dld*, *exaA*, *frdA*, *frdB*, *fumA*, *ldh*, *lldD*, *mqo*, *poxB*, *ppc*, and *pta*). A custom reference dataset was used to annotate the N_2_O reductase *nosZ* clade I and II genes. This dataset was updated from the one previously described by Sennett et al. [16] to include 45-full length reference sequences per gene, based on the phylogenetic tree in Supplementary Figure S2 of Conthe et al. [17]. All 90 sequence references are listed in Supplementary Table S2. Reads were also aligned to the *onr* gene that encodes the octaheme nitrite reductase (ONR), a close homolog of NrfA [18] using the custom dataset containing 22 sequences described in Saghaï et al. [19].

The DIAMOND outputs were converted to m8 blast format and downstream filtering and final assignments were performed in RStudio using the custom code described in detail in Sennett et al. 2024. Output of matching reads were normalized to reads per million of total reads (RPM) to account for differing sequencing depths. Reads derived from specific genes that met the assigned quality cutoffs were extracted from read sets using filterbyname from BBTools suite of programs [3]. The extracted reads were then uploaded to KBase and taxonomic assignment was performed using KAIJU (v1.9.0) [4] using default settings.

**Supplementary Tables**

**Supplementary Table S1. Calculation of added nitrate converted to ammonium via dissimilatory nitrate reduction to ammonium (DNRA) using isotopic measurements of ^15^N in a soil-extracted bacterial community with a clover extract as a carbon source under denitrifying conditions.** Experimental vials received an intial concentration of 2mM 6.3 At% ^15^N- labelled NO_3_^-^ (100 µmol NO_3_^-^ vial^-1^). These vials were then analyzed at the end of the incubation for ^15^N-NH_4_^+^ after chemical conversion of NH_4_^+^ to N_2_O using the sodium azide method (NaN_3_). ^15^N abundance was analyzed using a Finnigan Delta Plus XP isotope ratio mass spectrometer (IRMS) coupled to a PreCon. At% of ^15^N was corrected for scale and drift using in-house standards which had been calibrated to the international standards USGS25 and USGS26. To calculate excess ^15^N At% above natural abundance, the At% of unlabeled negative control samples was subtracted from the At% of the labeled samples. Data represents the mean value (n = 5 biological replicates).

**NH_4_^+^ concentration at the end of incubation**

559.5 ± 19.7 µmol vial^-1^

**Excess ^15^N At% in the NH_4_^+^ pool at the end of incubation (via the chemical conversion to N_2_O)**

0.37% ± 0.014

**Excess ^15^N At% in added NO_3_^-^ solution**

5.93% [6.3% – ^15^N natural At% of 0.367%]

**^15^N-NO_3_^-^ enrichment at the start of the incubation**

5.9 µmol vial^-1^

**^15^N-NH_4_^+^ enrichment at the end of incubation**

2.072 µmol vial^-1^

**% added NO_3_^-^ converted to NH_4_^+^ via DNRA**

35%

**Supplementary Table S2.** Information on NosZ protein dataset used for clade I/clade II classification.

| Organism (Conthe et al. 2018) | Protein Accession | NosZ Clade (Conthe et al. 2018) |
| --- | --- | --- |
| Pseudomonas aeruginosa PAO581 | AGV60993.1 | I |
| Burkholderia pseudomallei MSHR305 | AGR71407.1 | I |
| Burkholderia thailandensis H0587 | AHI65389.1 | I |
| Brucella abortus | AIJ50963.1 | I |
| Brucella suis | AIJ66148.1 | I |
| Methylophaga nitratireducenticrescens | AJO16109.1 | I |
| Pseudomonas fluorescens | AMZ70880.1 | I |
| Neisseria lactamica | ARB04292.1 | I |
| Neisseria mucosa | ARC51010.1 | I |
| Alcaligenes faecalis | ASC90844.1 | I |
| Sinorhizobium meliloti | ASJ61682.1 | I |
| Simonsiella muelleri ATCC 29453 | AUX61189.1 | I |
| Achromobacter xylosoxidans | AUZ18630.1 | I |
| Roseobacter denitrificans | AVL54235.1 | I |
| Rhodopseudomonas palustris | AVT76133.1 | I |
| Bradyrhizobium diazoefficiens | AWO87404.1 | I |
| Ralstonia solanacearum | AXV72269.1 | I |
| Paracoccus yeei | AYF00664.1 | I |
| Ralstonia pickettii | KFL22827.1 | I |
| Pseudomonas veronii | OPK05654.1 | I |
| Alcanivorax sp. | PHS58940.1 | I |
| Janthinobacterium sp. BJB312 | PHV31271.1 | I |
| Rhizobium sullae | PKA43318.1 | I |
| Photobacterium profundum | PSV62243.1 | I |
| Thauera sp. | PZU55555.1 | I |
| Paracoccus pantotrophus | RDD96771.1 | I |
| Azospirillum brasilense | RIW00398.1 | I |
| Rhizobium leguminosarum | WP_003590820.1 | I |
| Neisseria cinerea | WP_003675787.1 | I |
| Kingella denitrificans | WP_003784611.1 | I |
| Kingella kingae | WP_003787274.1 | I |
| Cardiobacterium hominis | WP_004143012.1 | I |
| Vibrio orientalis | WP_004414975.1 | I |
| Vibrio tubiashii | WP_004747054.1 | I |
| Aeromonas media | WP_005324393.1 | I |
| Lautropia mirabilis | WP_005675136.1 | I |
| Acidovorax delafieldii | WP_005797511.1 | I |
| Salinisphaera shabanensis | WP_006915220.1 | I |
| Cardiobacterium valvarum | WP_006984058.1 | I |
| Rhodanobacter fulvus | WP_007079923.1 | I |
| Oceaniovalibus guishaninsula | WP_007426212.1 | I |
| Rhodanobacter denitrificans | WP_007507670.1 | I |
| Rhodanobacter spathiphylli | WP_007806446.1 | I |
| Reinekea blandensis | WP_008045430.1 | I |
| Achromobacter arsenitoxydans | WP_008166018.1 | I |
| Azonexus hydrophilus | WP_028994793.1 | II |
| Campylobacter hyointestinalis | WP_059426052.1 | II |
| Campylobacter concisus | WP_103633399.1 | II |
| Mesoflavibacter zeaxanthinifaciens | WP_106677379.1 | II |
| Geobacillus thermodenitrificans | WP_099232746.1 | II |
| Solitalea longa | WP_103790248.1 | II |
| Arenibacter algicola | WP_093976926.1 | II |
| Riemerella anatipestifer | WP_064970925.1 | II |
| Myroides odoratimimus | WP_059050810.1 | II |
| Myroides injenensis | WP_050996164.1 | II |
| Runella slithyformis | WP_041339845.1 | II |
| Fulvivirga imtechensis | WP_040496853.1 | II |
| Mangrovimonas yunxiaonensis | WP_036119190.1 | II |
| Anditalea andensis | WP_035068779.1 | II |
| Salinimicrobium xinjiangense | WP_029036203.1 | II |
| Psychroserpens burtonensis | WP_028873327.1 | II |
| Gelidibacter mesophilus | WP_027124739.1 | II |
| Arenibacter certesii | WP_026815198.1 | II |
| Flavobacterium sasangense | WP_026726061.1 | II |
| Prevotella denticola | WP_025067032.1 | II |
| Flavobacterium saliperosum | WP_023575269.1 | II |
| Flavobacterium enshiense | WP_023573525.1 | II |
| Leptospira meyeri | WP_020775969.1 | II |
| Segetibacter koreensis | WP_018617447.1 | II |
| Nafulsella turpanensis | WP_017730956.1 | II |
| Pedobacter arcticus | WP_017259087.1 | II |
| Arcticibacter svalbardensis | WP_016196695.1 | II |
| Leptospira wolbachii | WP_015680928.1 | II |
| Desulfosporosinus meridiei | WP_014904216.1 | II |
| Desulforamulus ruminis | WP_013841561.1 | II |
| Nitratifractor salsuginis | WP_013553461.1 | II |
| Denitrovibrio acetiphilus | WP_013010250.1 | II |
| Sphaerobacter thermophilus | WP_012871219.1 | II |
| Imtechella halotolerans | WP_008236988.1 | II |
| Cellulophaga algicola | WP_041558227.1 | II |
| Leptospira biflexa | WP_012387413.1 | II |
| Bacillus massilionigeriensis | WP_075983258.1 | II |
| Hippea jasoniae | WP_051904383.1 | II |
| Melioribacter roseus | WP_041355904.1 | II |
| Altibacter lentus | WP_034258818.1 | II |
| Gaetbulibacter saemankumensis | WP_027137829.1 | II |
| Algoriphagus mannitolivorans | WP_026951434.1 | II |
| Pontibacter actiniarum | WP_025605765.1 | II |
| Flexithrix dorotheae | WP_020532851.1 | II |
| Desulfomonile tiedjei | WP_014808539.1 | II |

**Supplementary Figures**

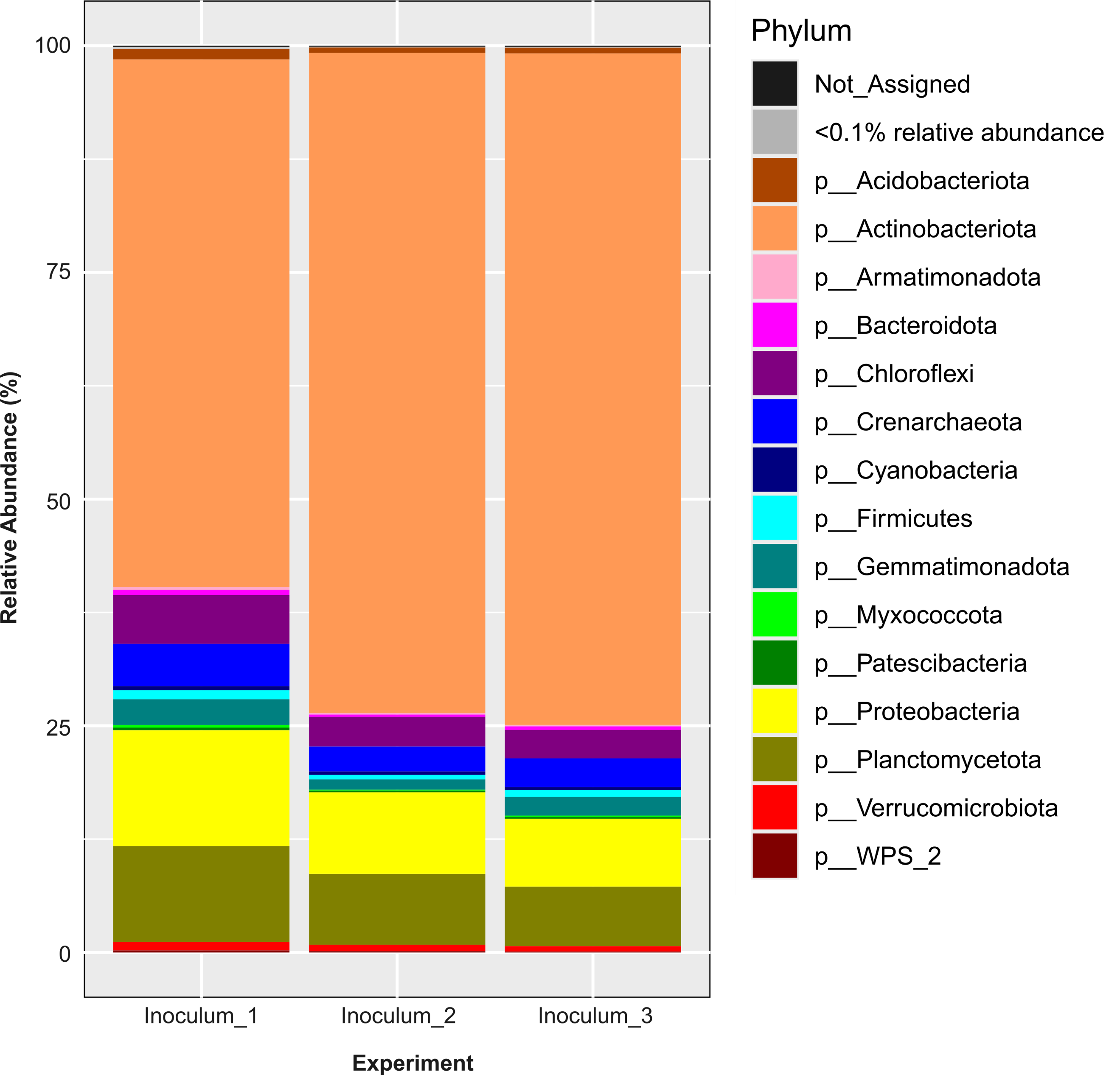

**Supplementary Figure S1. 16S rRNA gene-based taxonomic composition of soil-extracted bacterial communities used as inocula in Experiments 1, 2, and 3.** Inocula 1, 2, and 3 correspond to Experiments 1, 2, and 3, respectively. Taxonomic assignments are shown at the phylum level as relative abundances (%). Operational taxonomic units (OTUs) were clustered at a 97% sequence similarity threshold. Data represent mean values (n = 3).

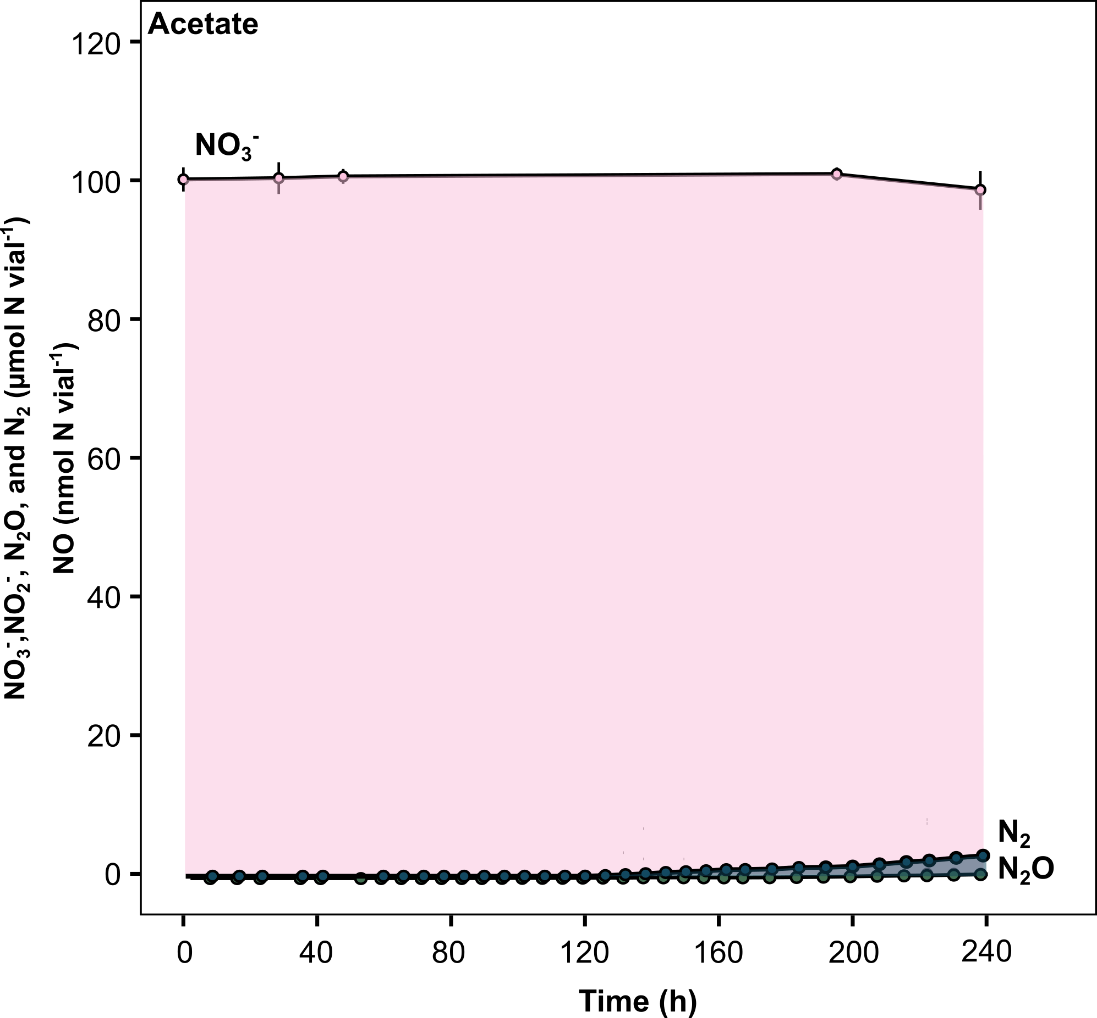

**Supplementary Figure S2. Denitrification phenotype of a soil-extracted bacterial community incubated with acetate as a carbon substrate.** Acetate was added at a rate of 0.015 g C vial^-1^ (0.3 g C L^-1^). The experimental vials had an initial cell concentration of 1.33 x10^9^ cells vial^-1^ and contained an initial concentration of 2 mM KNO_3_. The vials were made anoxic prior to the incubation. The NO concentration was variable, ranging between 0-75 nmol vial^-1^ throughout the incubation. Data are presented as mean values ± the standard deviation (*n* = 3 biological replicates).

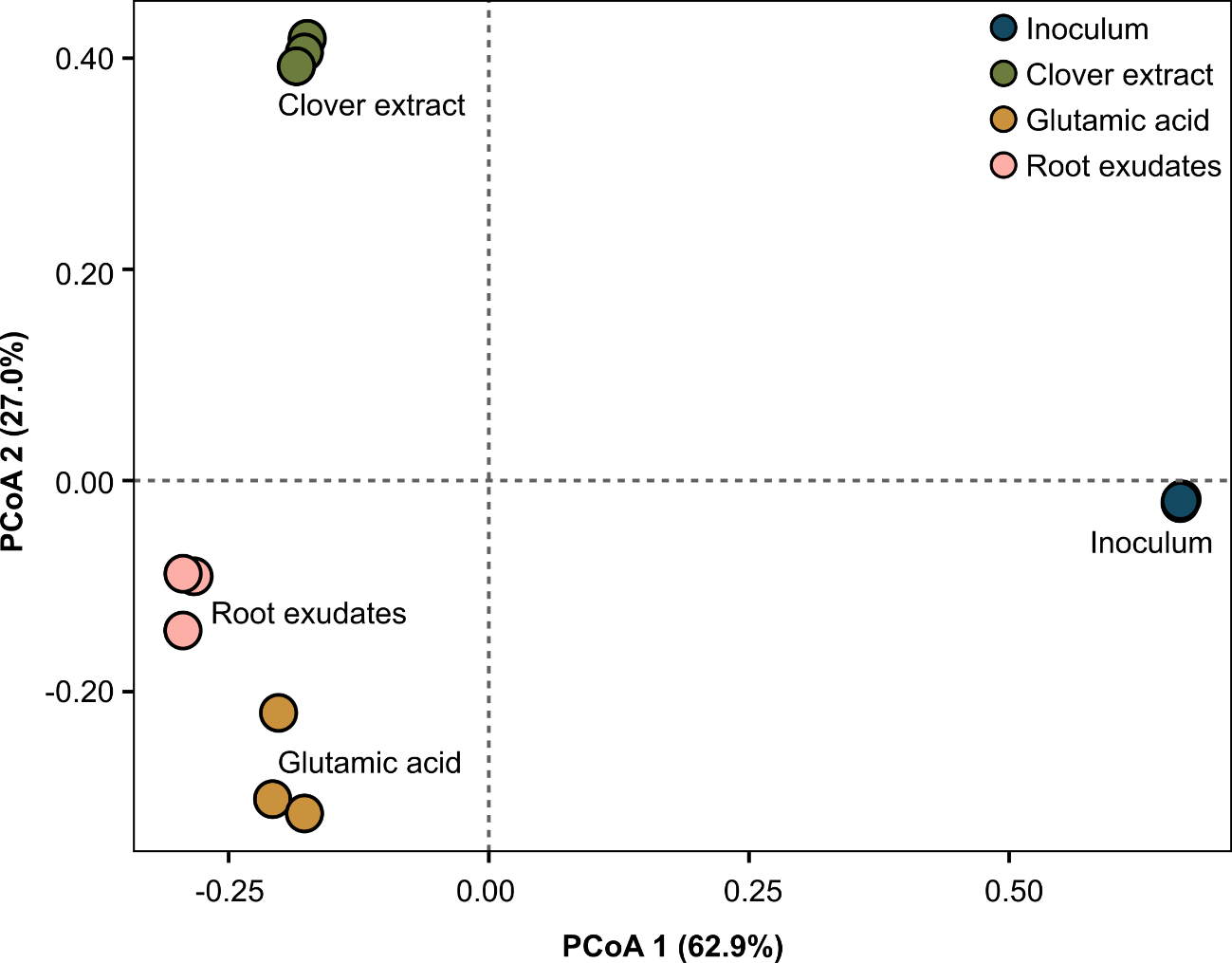

**Supplementary Figure S3. Beta-diversity of a soil-extracted bacterial community incubated with different carbon substrates under denitrifying conditions.** Principal coordinates analysis (PCoA) of a bray-curtis dissimilaritity matrix generated using operational taxonomic units (OTUs) clustered using a 97% similarity threshold. Treatments included the original inoculum and the bacterial community at the completion of denitrification using either a clover extract, glutamic acid, or an artificial root exudate cocktail as a carbon substrate. Individual biological replicates (*n* = 3) are presented.

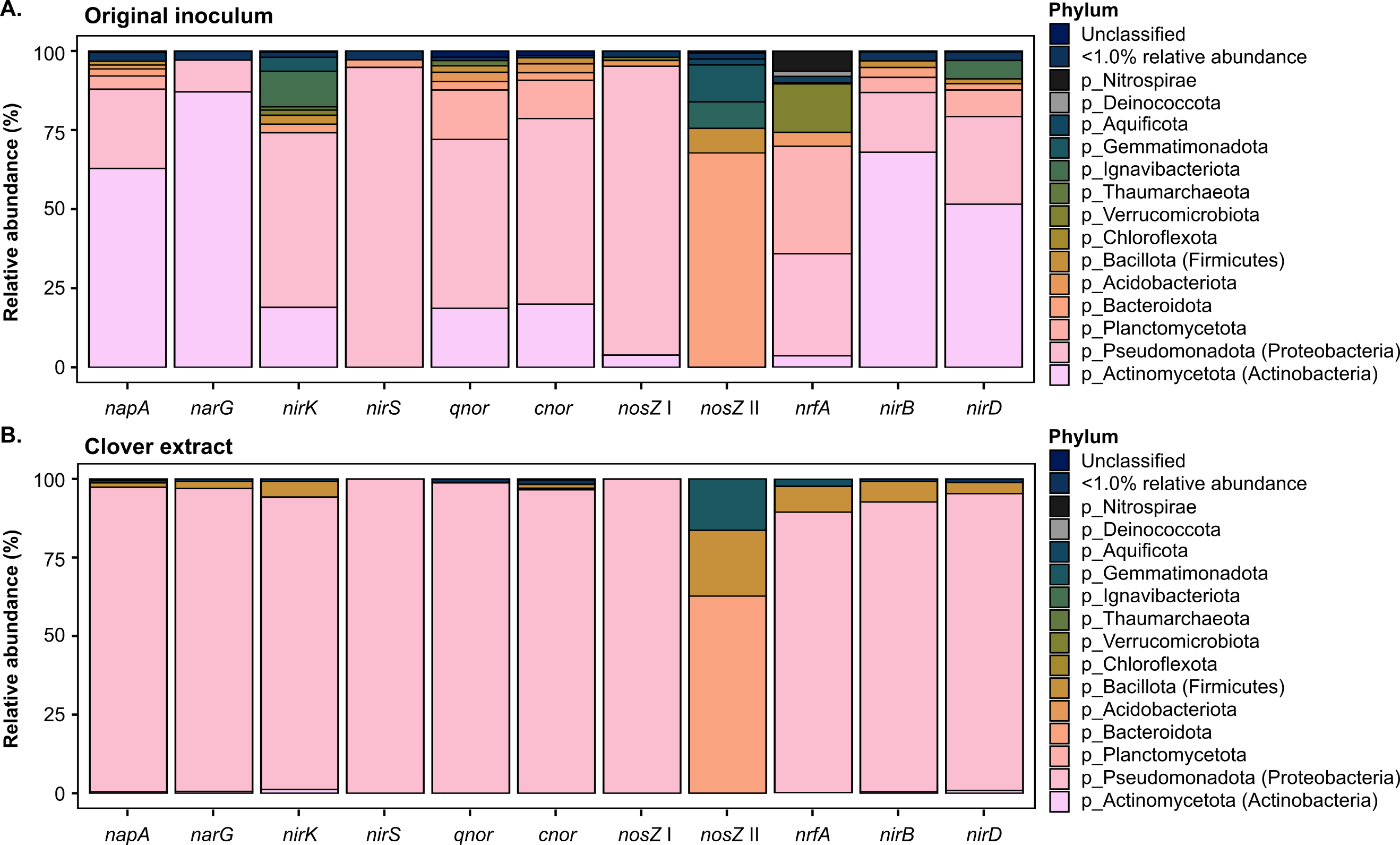

**Supplementary Figure S4. Taxonomic assignment at the phylum-level of reads encoding denitrification and DNRA genes from the metagenomes of a soil-extracted bacterial community incubated with clover extract as a carbon substrate under denitrifying conditions.** Metagenomes of the bacterial community from the **(a)** original inoculum and **(b)** clover extract treatment at the completion of denitrification. Reads encode NO_3_^-^ reductases (*napA* and *narG*), NO_2_^-^ reductases (*nirK*, *nirS*, *nrfA*, *nirB*, and *nirD*), NO reductases (*cnor* and *qnor*), and N_2_O reductases (*nosZ* clade I and II). The data represents the relative abundance presented as the mean values (*n* = 3 biological replicates).

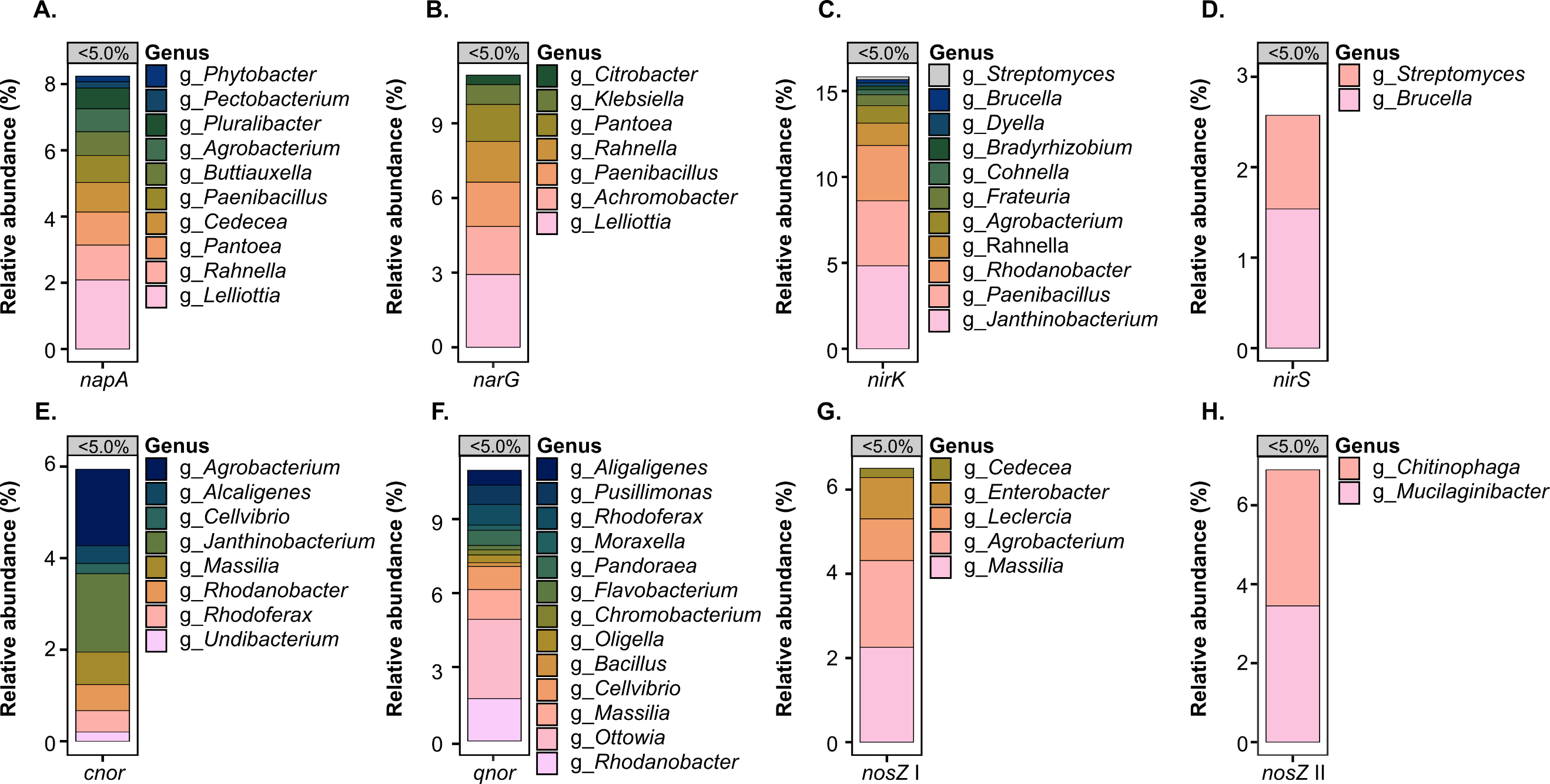

**Supplementary Figure S5.** **Taxonomic assignment at the genus-level of low abundance (<5% relative abundance) reads encoding denitrification genes from the metagenomes of a soil-extracted bacterial community incubated with clover extract as a carbon substrate under denitrifying conditions.** The metagenomes are of the bacterial community at the completion of denitrification. Reads encode NO_3_^-^ reductases *napA* and *narG* **(a, b)**, NO_2_^-^ reductases *nirK* and *nirS* **(c, d)**, NO reductases *cnor* and *qnor* **(e, f)**, and N_2_O reductases *nosZ* clade I and II **(g, h)**. The data represents the relative abundance presented as the mean values (*n* = 3 biological replicates).

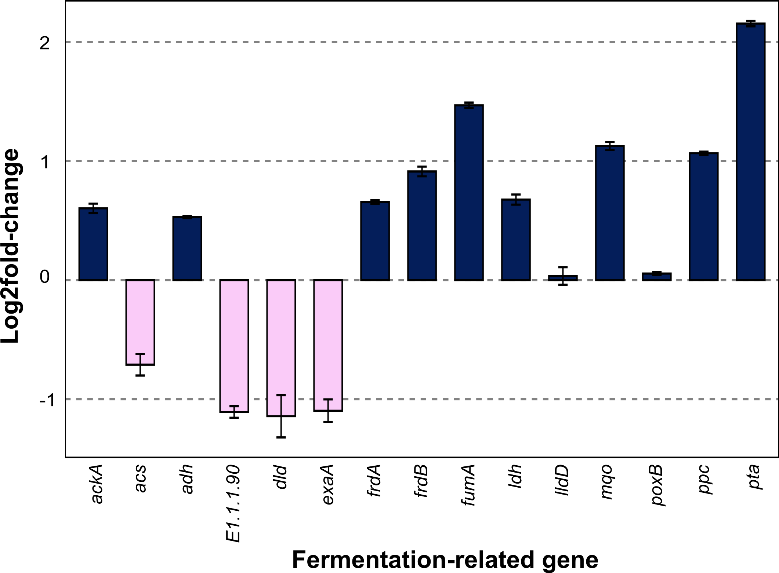

**Supplementary Figure S6. Log2fold-changes in the read abundance of fermentation-related genes in a soil-extracted bacterial community incubated with clover extract as a carbon substrate under denitrifying conditions.** Fold-changes were calculated by comparing gene abundance in the clover extract treatment at the completion of denitrification to the original inoculum. Fermentation-related genes include: acetate kinase (*ackA*), acetyl-CoA synthetase (*acs*), alcohol dehydrogenase (*adh*), aryl-alcohol dehydrogenase (*E.1.1.90*), D-lactate dehydrogenase (*dld*), alcohol dehydrogenase (*exaA*), fumarate reductase flavoprotein subunit (*frdA*), fumarate reductase iron-sulfur subunit (*frdB*), fumarate hydratase (*fumA*), L-lactate dehydrogenase cytochrome (*lldD*), malate dehydrogenase quinone (*mqo*), pyruvate dehydrogenase (*poxB*), phosphoenolpyruvate carboxylase (*ppc*), and phosphate acetyltransferase (*pta*). Reads were normalized to reads per total million reads prior to analysis. Data presented as the mean values ± the standard deviation (*n* = 3 biological replicates).

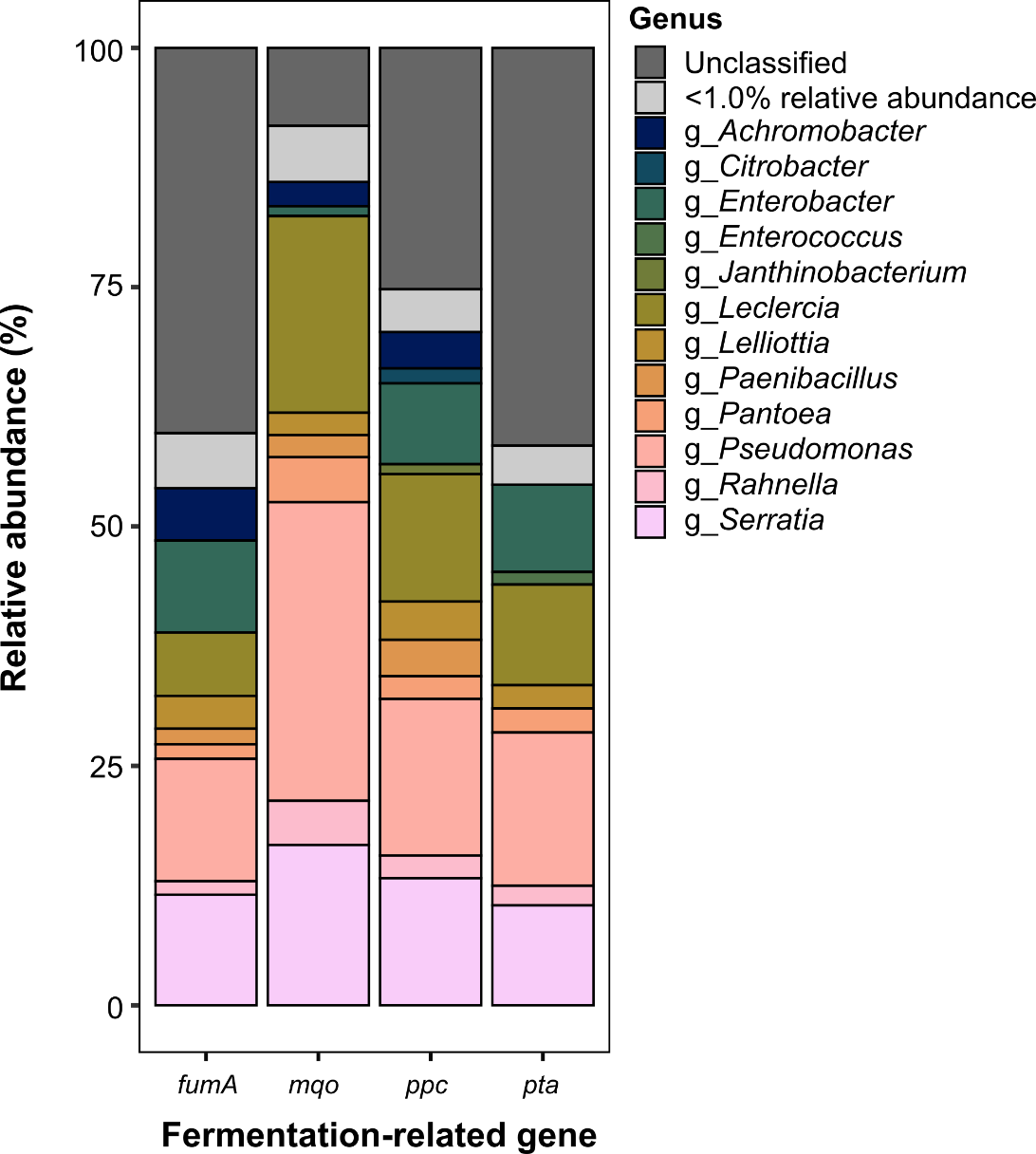

**Supplementary Figure S7. Taxonomy assignment of reads encoding select fermentation-related genes from the metagenomes of a soil-extracted bacterial community incubated with clover extract as a carbon substrate under denitrifying conditions.** Metagenomes are of the bacterial community at the completion of denitrification. The select fermentation-related genes include fumarate hydratase (*fumA*), malate dehydrogenase quinone (*mqo*), phosphoenolpyruvate carboxylase (*ppc*), and phosphate acetyltransferase (*pta*). The data represents the relative abundance presented as the mean values (*n* = 3 biological replicates).

**References**

1.	Østby H, *et al.* Chromatographic analysis of oxidized cello-oligomers generated by lytic polysaccharide monooxygenases using dual electrolytic eluent generation. *Journal of Chromatography A* 2022;**1662**:462691. https://doi.org/0.1016/j.chroma.2021.462691

2.	Liu B, Frostegård Å, Bakken LR. Impaired reduction of N_2_O to N_2_ in acid soils is due to a posttranscriptional interference with the expression of *nosz*. *mBio* 2014;**5**:e01383-14. https://doi.org/10.1128/mBio.01383-14.

3.	Molstad L, Dörsch P, Bakken LR. Robotized incubation system for monitoring gases (O_2_, NO, N_2_O, N_2_) in denitrifying cultures. *J Microbiol Methods* 2007;**71**:202–11. https://doi.org/10.1016/j.mimet.2007.08.011.

4.	Molstad L, Dörsch P, Bakken LR. Improved robotized incubation system for gas kinetics in batch cultures. *Researchgate*. 2016. https://doi.org/10.13140/RG.2.2.30688.07680.

5.	Lim NYN, Frostegård Å, Bakken LR. Nitrite kinetics during anoxia: The role of abiotic reactions versus microbial reduction. *Soil Biol Biochem* 2018;**119**:203–09. https://doi.org/https://doi.org/10.1016/j.soilbio.2018.01.006

6.	Bolyen E, *et* *al*. Reproducible, interactive, scalable and extensible microbiome data science using QIIME 2. *Nature Biotechnology* 2019;**37**:852-57. https://doi.org/https://doi.org/10.1038/s41587-019-0190-3.

7.	Callahan B, McMurdie P, Rosen M *et al.* Dada2: High-resolution sample inference from illumina amplicon data. *Nat Methods* 2016;**13**: 581–83. https://doi.org/https://doi.org/10.1038/nmeth.3869

8.	Quast C, *et al.* The silva ribosomal rna gene database project: Improved data processing and web-based tools. *Nucleic Acids Res* 2013;**41**:590-96. https://doi.org/doi:10.1093/nar/gks1219.

9.	Dhariwal A, *et al.* Microbiomeanalyst: A web-based tool for comprehensive statistical, visual and meta-analysis of microbiome data. *Nucleic Acids Res* 2017;**45**:180-88. https://doi.org/10.1093/nar/gkx295.

10.	Chong J, *et al.* Using microbiomeanalyst for comprehensive statistical, functional, and meta-analysis of microbiome data. *Nat Protoc* 2020;**15**:799-821. https://doi.org/10.1038/s41596-019-0264-1

11.	Arkin AP, *et al.* KBase: The united states department of energy systems biology knowledgebase. *Nat Biotechnol* 2018;**36**:566–69. https://doi.org/https://doi.org/10.1038/nbt.4163

12.	Bolger AM, Lohse M, Usadel B. Trimmomatic: A flexible trimmer for illumina sequence data. *Bioinformatics* 2014;**30**:2114-20. https://doi.org/10.1093/bioinformatics/btu170

13.	Buchfink B, Xie C, Huson DH. Fast and sensitive protein alignment using diamond. *Nature Methods* 2015;**12**:59-60. https://doi.org/10.1038/nmeth.3176

14.	Mendler K, *et al.* Annotree: Visualization and exploration of a functionally annotated microbial tree of life. *Nucleic Acids Research* 2019;**47**:4442-48. https://doi.org/10.1093/nar/gkz246

15.	Pandey CB, *et al.* DNRA: A short-circuit in biological N-cycling to conserve nitrogen in terrestrial ecosystems. *Sci Total Environ* 2020;**788**:139710. https://doi.org/https://doi.org/10.1016/j.scitotenv.2020.139710

16.	Sennett LB, *et al.* Determining how oxygen legacy affects trajectories of soil denitrifier community dynamics and N_2_O emissions. *Nature Communications* 2024;**15** https://doi.org/https://doi.org/10.1038/s41467-024-51688-w

17.	Conthe M, *et al.* Life on N_2_O: Deciphering the ecophysiology of N_2_O respiring bacterial communities in a continuous culture. *ISME J* 2018;**12**:1142-53.

18.	Tikhonova TV, Trofimov AA, Popov VO. Octaheme nitrite reductases: Structure and properties. *Biochemistry (Moscow)* 2012;**77**:1129-38. https://doi.org/10.1134/S0006297912100057

19.	Saghaï A, *et al.* Phyloecology of nitrate ammonifiers and their importance relative to denitrifiers in global terrestrial biomes. *Nat Commun* 2023;**14**:8249. https://doi.org/https://doi.org/10.1038/s41467-023-44022-3
